# Structural variation, selection, and diversification of the *NPIP* gene family from the human pangenome

**DOI:** 10.1101/2025.02.04.636496

**Authors:** Philip C. Dishuck, Katherine M. Munson, Alexandra P. Lewis, Max L. Dougherty, Jason G. Underwood, William T. Harvey, PingHsun Hsieh, Tomi Pastinen, Evan E. Eichler

## Abstract

The *NPIP* (nuclear pore interacting protein) gene family has expanded to high copy number in humans and African apes where it has been subject to an excess of amino acid replacement consistent with positive selection (1). Due to the limitations of short-read sequencing, *NPIP* human genetic diversity has been poorly understood. Using highly accurate assemblies generated from long-read sequencing as part of the human pangenome, we completely characterize 169 human haplotypes (4,665 *NPIP* paralogs and alleles). Of the 28 *NPIP* paralogs, just three (*NPIPB2*, *B11*, and *B14*) are fixed at a single copy, and only a single locus, *B2*, shows no structural variation. Four *NPIP* paralogs map to large segmental duplication blocks that mediate polymorphic inversions (355 kbp–1.6 Mbp) corresponding to microdeletions associated with developmental delay and autism. Haplotype-based tests of positive selection and selective sweeps identify two paralogs, *B9* and *B15*, within the top percentile for both tests. Using full-length cDNA data from 101 tissue/cell types, we construct paralog-specific gene models and show that 56% (31/55 most abundant isoforms) have not been previously described in RefSeq. We define six distinct translation start sites and other protein structural features that distinguish paralogs, including a variable number tandem repeat that encodes a beta helix of variable size that emerged ∼3.1 million years ago in human evolution. Among the 28 *NPIP* paralogs, we identify distinct tissue and developmental patterns of expression with only a few maintaining the ancestral testis-enriched expression. A subset of paralogs (*NPIPA1*, *A5*, *A6-9*, *B3-5*, and *B12/B13*) show increased brain expression. Our results suggest ongoing positive selection in the human population and rapid diversification of *NPIP* gene models.

## INTRODUCTION

*NPIP* (also known as *morpheus*) is a gene family of unknown function that has undergone independent duplication in several primate lineages (1–3). The gene family was first described based on the observation of a rapid expansion in African apes where the underlying genes show a significant excess of amino acid replacements (extraordinary dN/dS values) consistent with the action of positive selection (1). The ∼20 kbp duplicon that contains *NPIP*, LCR16a, is interspersed across human chromosome 16 (Fig. 1A) (4, 5) with a solitary copy on human chromosome 18 (1). These LCR16 duplications mediate recurrent duplications and deletions frequently associated with neurodevelopmental delay (6–8), including one of the most common genetic causes of autism spectrum disorder (9, 10). Altogether, the segmental duplications (SDs) associated with LCR16a (2) span ∼10% of the euchromatic portion of human chromosome 16p, having emerged and expanded since ape divergence from the Old World monkeys (25 million years ago [mya]). The LCR16a-encoding *NPIP* has been described as a “core duplicon” for its characteristic overabundance within these intrachromosomal duplications (11). It has independently duplicated at least five times over the course of primate evolution leading each time to the formation of interspersed SDs where lineage-specific duplications accrue flanking the core LCR16a duplicon (2, 3). Although the gene model has significantly changed among primates, the open reading frame (ORF) has been maintained over the course of primate evolution (3).

**Figure 1.**
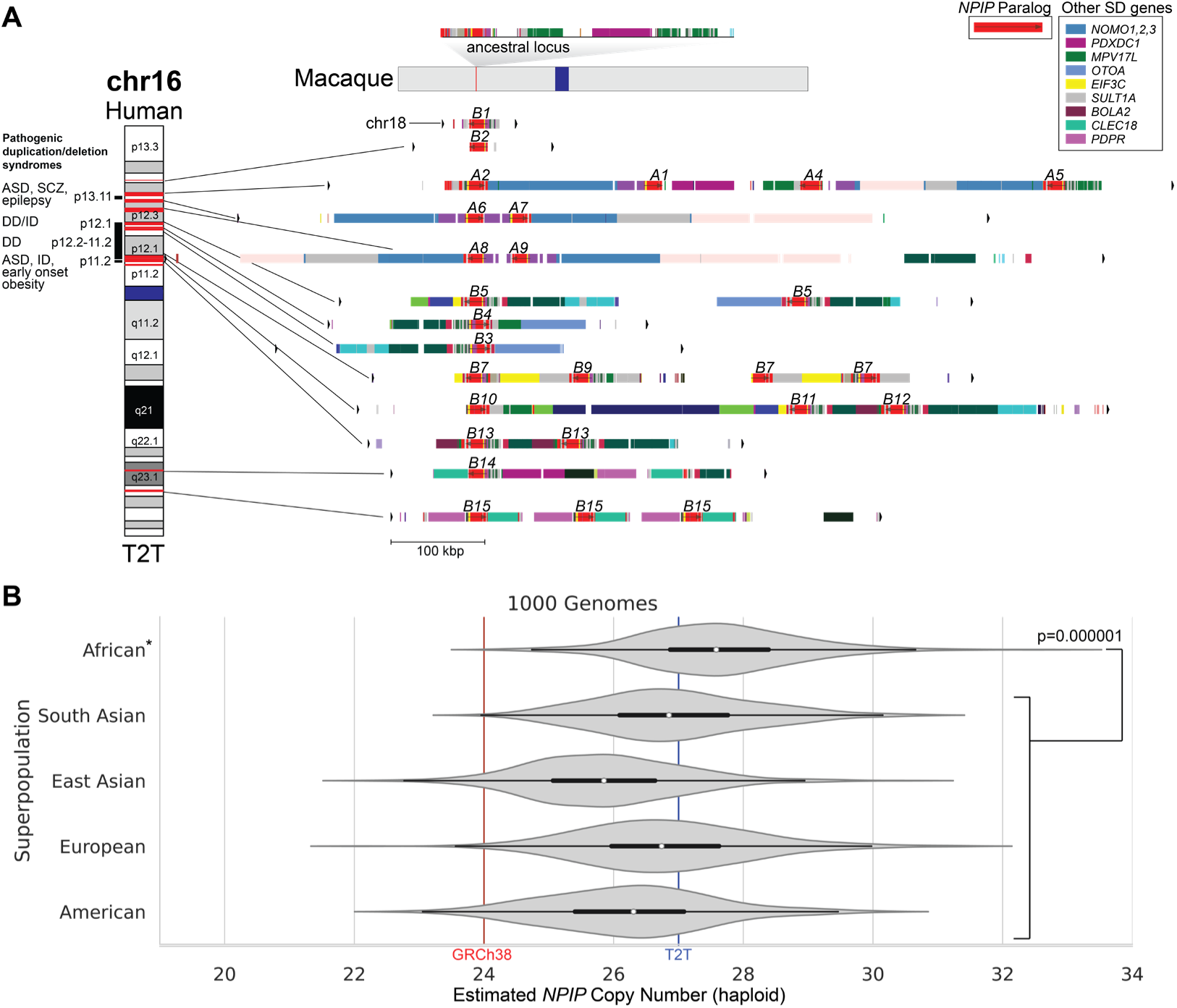
*NPIP* locus organization and copy number variation. **A)** *NPIP* regional organization in the T2T-CHM13 genome. The single-copy sequence in the macaque genome (Mmul10; top) is compared to the duplicated sequence on human chromosomes 16 and 18. Red highlights on chromosome 16 (left) correspond to the *LCR16a* duplicon encoding different *NPIP* genes, with segmental duplication (SD) content annotated by DupMasker (colored bars) (97). *NPIP* gene names (*A1-9*, *B1-15*) are labeled above DupMasker tracks, with the colored bars indicating other chromosome 16 genes (key) associated with the *NPIP* expansion in humans. To the left of the ideogram, pathogenic duplication/deletion syndromes associated with *NPIP* SDs are shown as red horizontal bars on the ideogram (ASD=autism spectrum disorder, SCZ=schizophrenia, DD=developmental delay, ID=intellectual disability). These recurrent deletions and duplications are mediated by *NPIP-*containing SD blocks. **B)** Read-depth estimates (fastCN) of modern human per-haplotype *NPIP* copy number from the 1000 Genomes Project (n=2,609), grouped by superpopulation.

Because *NPIP* is frequently embedded in large blocks of SDs that share >97% sequence identity (Fig. 1), standard sequencing and assembly methods have limited our understanding of its genetic diversity and, consequently, our ability to make genetic associations or perform standard population genetic analyses. FISH analysis with LCR16a probes along with read-depth analyses using short reads, however, have been used to estimate a range of 20-30 copies per human haplotype (2, 3). Targeted bacterial artificial chromosome (BAC) assemblies have partially resolved the *NPIP* loci in the most commonly used reference genomes. Because of the high sequence identity of the underlying duplicated segments, misassembly of the loci has been frequently encountered over the last 20 years of human genome assemblies. Even in one of the most recent human genome references, GRCh38, there is evidence of at least two chimeric misassemblies created due to inadvertently assembling paralogous loci whose sequence identity approximates allelic variation. Not surprisingly, GRCh38 contains just 24 copies of LCR16a, compared to the median 25 copies estimated by short-read WGS read depth to be present in most human haplotypes (12).

Over the last few years, a series of resources and methods have been developed making it possible to systematically characterize human genetic variation and expression across these regions of chromosome 16, arguably for the first time. First, the T2T (Telomere-to-Telomere) Consortium recently completed the assembly of a single human haplotype, CHM13, by combining highly accurate, long HiFi (high-fidelity) reads with ultra-long ONT reads (13). As a result, all *NPIP* gene copies are fully resolved in this haplotype providing a complete reference for comparisons. Second, both the HPRC (Human Pangenome Reference Consortium) and HGSVC (Human Genome Structural Variation Consortium) using similar approaches have published and released contiguous phased assemblies of 80 unrelated individuals (14–16). The availability of both short-read sequencing and long-read sequencing, including phasing information, enables the characterization and validation of entire chromosomal haplotypes for even the most identical gene families (15, 17, 18). Third, the recent advancements of multiple sequence alignment (MSA) and phylogenetic methods optimized for comparing thousands of viral genomes (19, 20) have facilitated the evolutionary reconstruction of rapidly evolving 20 kbp segments of human DNA like LCR16a. We directly apply these methods to characterize thousands of *NPIP* paralogs and alleles to reconstruct the complex population genetic history underlying these regions of human chromosome 16, including the mutational forces that have shaped them. Finally, the recent release of 1.4 billion full-length cDNA from 384 Iso-Seq libraries from the Genomic Answer for Kids Study and ENCODE, among others (21–35), makes it possible to assign transcript data to specific paralogs and alleles—a near impossibility previously with traditional short-read RNA-seq data. We use these data to accurately construct gene models, define transcription start sites, distinguish potential protein-coding genes from pseudogenes, and interrogate expression and population genetic properties for specific *NPIP* copies.

## RESULTS

### Human genetic diversity

Using the complete sequence of the T2T human genome assembly, we first annotated LCR16a and its associated SDs using DupMasker for the T2T-CHM13 reference genome (Fig. 1A). The analysis reveals 27 *NPIP* genes—26 of which map to 12 duplication blocks on chromosome 16 and a solitary copy mapping to chromosome 18, as expected (Johnson et al, 2001). This is in stark contrast to the macaque genome (Fig. 1A) where only a single copy of LCR16a was identified. We assigned gene names based on best matches according to GRCh38 gene annotation. To estimate the copy number distribution in the human population, we mapped whole-genome shotgun sequence (WGS) data (36) from the 2,609 unrelated individuals from the 1000 Genomes Project (1KG) using read depth to estimate the diploid and haploid copy number across each superpopulation. Among humans, we estimate the haploid copy number ranges from 21 to 33 copies with the highest copy number observed among individuals of African descent (Wilcoxon rank-sum test, p=0.000001, Fig. 1B).

To understand the variation in structure of *NPIP* loci across the human population, we collected previously assembled and released genomes from the HPRC (n=43) and HGSVC (n=37) (14–16), along with a draft T2T assembly of HG002 (37), two individuals from Papua New Guinea (38), the reference genome T2T-CHM13 v2.0 (13), GRCh38, and an additional haploid cell line CHM1 (39, 40), for a total of 169 unrelated haplotypes (Supplementary Table S2). We identified and extracted *NPIP* loci from each assembly by aligning the *NPIP* locus from GRCh38 to each haplotype with minimap2 and wfmash (Methods), for a total of 4,665 copies of *NPIP.* As *NPIP* loci are known to be structurally variable and subject to gene conversion, we did not rely on synteny alone to determine the paralog identity. Instead, we created an MSA and maximum likelihood phylogeny of the 4,665 *NPIP* loci from the 169 assembled haplotypes, using Bornean orangutan (*Pongo pygmaeus*) and siamang (*Symphalangus syndactylus*) as outgroups (Fig. 2A). Monophyletic clades with >75% branch support (SH-aLRT) were used to assign copies to one of 28 defined paralogs, named based on phylogenetic identity to T2T-CHM13 and GRCh38. In cases where a clade did not have an anchor in T2T-CHM13 or GRCh38, we defined it based on its nearest neighbor (i.e., *NPIPA1L*). In cases where there was insufficient genetic distance to distinguish paralogs, they were grouped into a single clade comprising the two copies (i.e., *NPIPB12/B13*). For simplicity, we subsequently shorten gene names by dropping the *NPIP* prefix in this report. Additionally, to estimate the age of each branch, we created a timetree with LSD2 (Methods) (41), incorporating paralogs from human assemblies CHM13, CHM1, GRCh38, HG002, and PNG15, along with nonhuman primate sequences from the T2T Primate Project (*Pan troglodytes*, *Pan paniscus*, *Gorilla gorilla*, *Pongo pygmaeus*, *Pongo abelii*, and *Symphalangus syndactylus*) (42), and the single-copy ancestral *NPIP* gene from *Macaca fascicularis* (43) as the outgroup (Fig. S1).

**Figure 2.**
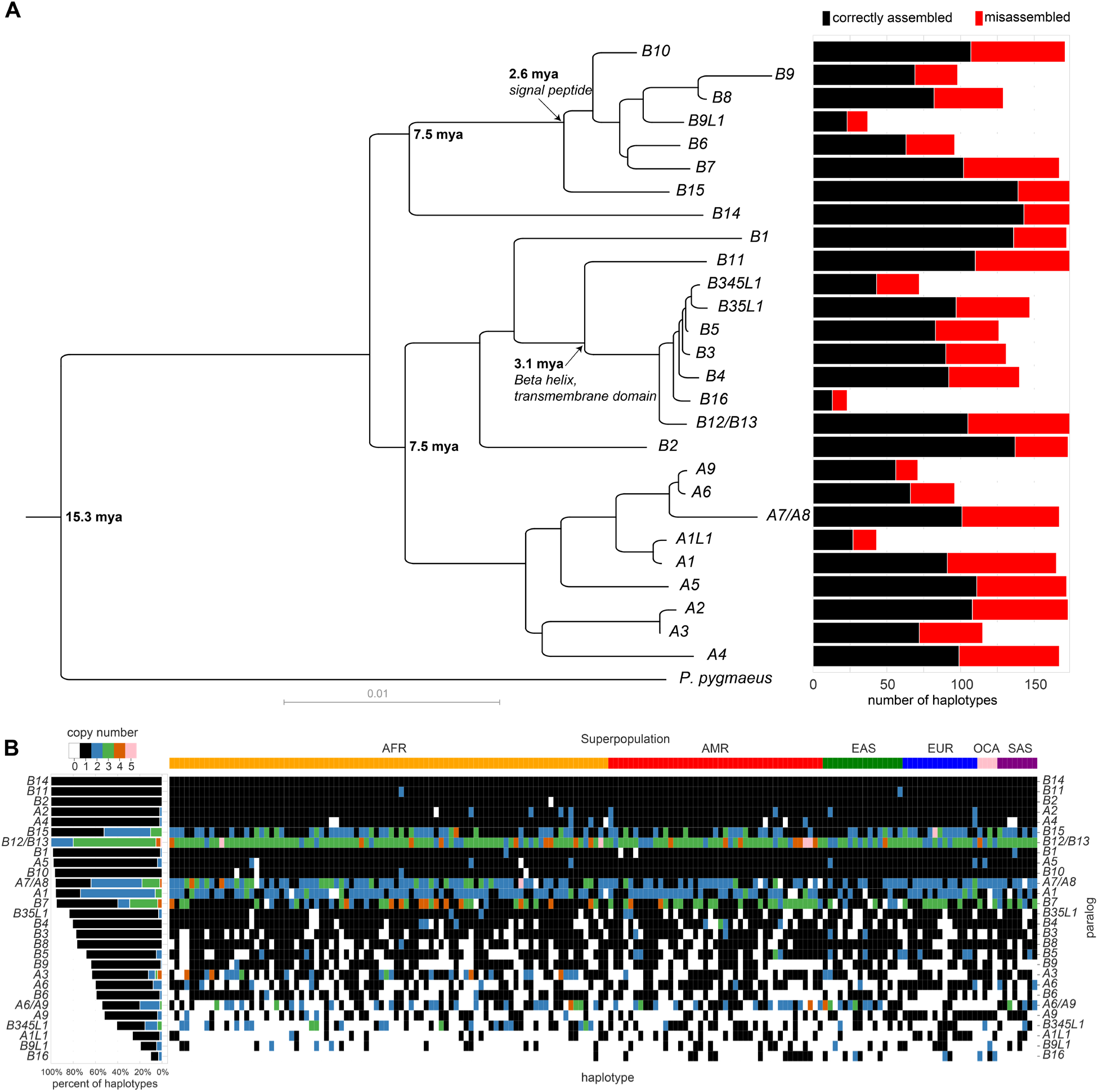
Classification of human *NPIP* haplotypes and locus-specific copy number. **A)** Left: Maximum likelihood phylogeny of human *NPIP* loci constructed using *Pongo pygmaeus* as an outgroup. It is based on 15 kbp of intronic (noncoding) sequence. Right: Frequency of each paralog among the 169 haplotypes passing assembly validation (black) and number of misassembled loci typically where a potential collapse was identified (red). **B)** Copy number summary of 169 assembled haplotypes. Color indicates copy number of each gene, as defined by the phylogenetic grouping (panel A). Paralogs are sorted by fraction of validated haplotypes containing at least one copy. Left: Percent of haplotypes with each copy number state for each paralog, restricted to assembled regions passing QC. Right: Copy number states for all assembled haplotypes, with each column representing a separate haplotype. Haplotypes are grouped by continental superpopulation (above) (AFR: Africa, AMR: the Americas, EAS: East Asia, EUR: Europe, OCA: Oceania, SAS: South Asia).

Even among long-read sequence-assembled genomes, high sequence identity duplications that are hundreds of kbp in length remain a common source of misassembly and collapse (44). We, therefore, validated the integrity of each assembled haplotype using computational tools designed to detect misassemblies (i.e., NucFreq, Flagger, and GAVISUNK (15, 39, 45)). For a haplotype structure to be classified as correctly assembled, we required contiguous assembly without collapse across all duplicated segments (not just *NPIP*) (Fig. 1A), including at least 30 kbp of flanking unique sequence. The assembly validation rate varied from 52-85% (Fig. 2A, right). The copy number of *NPIP* paralogs varies widely across assembled haplotypes (Fig. 2B). Copy number heterozygosity for the paralogs, defined as the frequency of discordant copy numbers between the two haplotypes of a sample, ranges from 0 for *B2*, *B11*, and *B14*, with all individuals having just one copy, to 0.74 for *A6/A9*. While only *B2*, *B11*, and *B14* are fixed in copy number, *A2*, *A4*, *B12/B13*, and *B15* always have at least one copy in all assemblies that pass QC. Individual members of *NPIP* subfamilies *B3-B5* and *B6-9* are not always present when considered individually, yet at least one paralog from each of these larger subfamilies is always present in a given haplotype (Fig. 2B).

In addition to copy number variation, interlocus gene conversion (IGC) is another common source of *NPIP* variation, as the high sequence identity among paralogs enables the replacement of *NPIP* sequence from one paralog to another. As a result, the sequence content of a paralog does not always correspond to the same syntenic location when compared to other human haplotypes. We reanalyzed a recent genome-wide IGC callset for a subset of these haplotypes (n=94) to classify IGC patterns among *NPIP* duplication blocks (46). As expected, IGC between paralogs is frequent (exceeding >50% of haplotypic configurations) and is driven primarily by proximity (1-2 Mbp) with six distinct IGC “hotspots” identified (Figs. S2, 3A) on the short arm of human chromosome 16. Active sites of gene conversion frequently correspond to the breakpoints associated with recurrent human microdeletion and microduplication syndromes (Fig. 3A). IGC occurs within but not between the two major subfamilies (*NPIPB* copies undergo IGC only with *NPIPB* but not *NPIPA* loci). We also observe particular biases in donor/acceptor directionality. For example, the putative ancestral paralog *NPIPA1* acts only as a donor to *A5*, *A6*, *A8*, and *A9* locations in distal chromosome 16p, but never as an acceptor, reflecting either functional constraint or bias in the mutation process itself.

**Figure 3.**
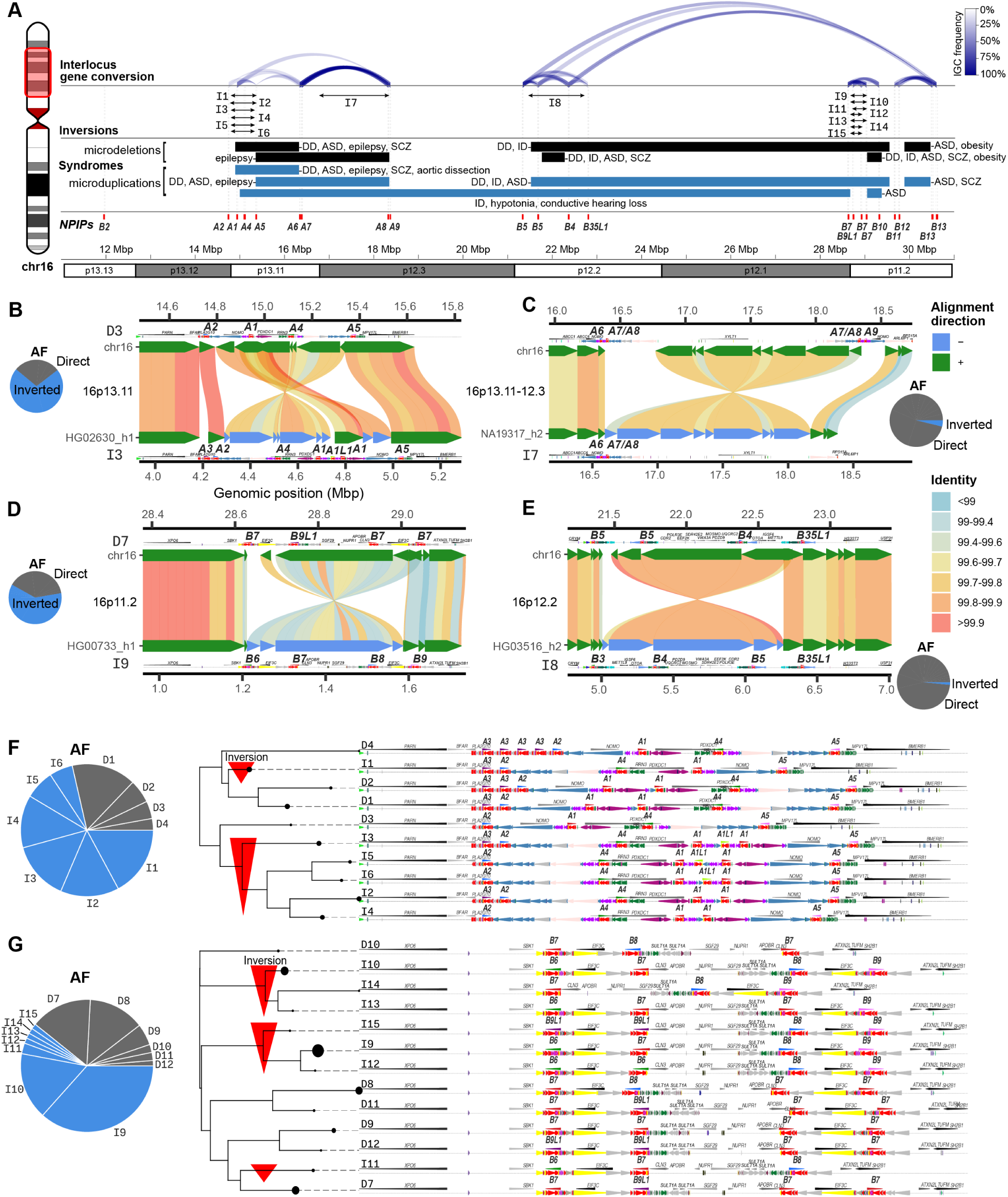
*NPIP* interlocus gene conversion (IGC) and complex structural changes. **A)** Overview of *NPIP* loci on chromosome 16p (highlighted ideogram region). The location of each T2T-CHM13 *NPIP* paralog is shown as a red vertical bar. The count and location of IGC between *NPIP* pairs is shown as blue arcs at the top, with opacity corresponding to number of observed haplotypes. Inversions mediated by *NPIP* are shown as black arrows, named corresponding to structures in panels B-G. Known pathogenic microdeletions and microdeletions with breakpoints at *NPIP* are shown as black and blue bars, respectively (6–10, 67–82). **B-E)** Large-scale inversion polymorphisms associated with *NPIP* loci (below) as compared to T2T-CHM13 v2.0 (top). Inversions are shown with SVbyEye, with DupMasker annotations for each haplotype. Allele frequency (AF) for inverted (I) and direct (D) orientation haplotypes are shown with the pie charts. **F-G)** The duplication architecture (DupMasker) of the *A1-5* and *B6-9* loci for the most common haplotype configurations, grouped by a neighbor-joining tree of double-cut-and-join edit distance (pairwise number of rearrangements between configurations). *NPIP* sequence denoted in red. The size of the circle for each clade corresponds to frequency of each configuration, and direct and inverted orientation configurations are named by frequency. Red arrows under the cladogram indicate configurations inverted with respect to T2T-CHM13.

During our comparative analysis, we frequently noted that the gene order of unique (nonduplicated) genes bracketed by *NPIP* copies are inverted in different human haplotypes. Across chromosome 16p, we identify four inversion polymorphisms ranging in size from 350 kbp to 1.6 Mbp in size (Fig. 3B-E). The breakpoints of these inversions map either at *NPIP* copies or associated SDs flanking *NPIP.* All of these large inversions are common polymorphisms (>5% allele frequency) and in some cases represent the major allelic configuration in the human population (47, 48). In several cases, the inverted unique sequence shows considerable allelic divergence (>0.2%) suggesting a deep coalescence as has been observed for other human inversion polymorphisms on other chromosomes (47, 49). Indeed, the coalescence of the D7 inversion polymorphism at 16p11.2 was previously estimated as 1.35 mya and associated with susceptibility to asthma and obesity (50, 51). González et al. estimated that at least six distinct haplotypes exist at this locus based on multidimensional scaling of single-nucleotide polymorphisms (SNPs). Using phased genomes, we double this number resolving 13 structural configurations at this locus distinguished by orientation, *NPIP* paralog identity, and *SULT1A* copy number that were previously indistinguishable. This complete sequence resolution may help explain their observed association of the inversion with increased *SULT1A4* expression and decreased *SULT1A1* expression (50). Notably, we find that many of the human haplotypic configurations occurred in conjunction with copy number variation and IGC events associated with specific *NPIP* loci.

To more systematically classify different structural configurations, we encode haplotypes by the identity, order, and orientation of *NPIP* paralogs and marker genes. We apply a double-cut-and-join rearrangement distance metric (Methods) (52) between each configuration to create corresponding neighbor-joining trees for each of the major *NPIP* clusters (Fig. 3E-F). For example, at the ancestral chromosome 16p13.11 locus, we observe a 545 kbp inversion and the variable presence or absence of *A3* and a newly discovered paralog, *A1L1.* By contrast, *A5* maps invariably at the proximal end of this cluster (Fig. 3A,E). The *A1L1* paralog only associates with 16p13.11 haplotypes that are inverted relative to T2T-CHM13; this 545 kbp inversion is the major allele (AF=0.69). The chromosome 16p11.2 locus contains *NPIPB6*, *B7*, *B8*, and *B9* spanning a 650 kbp inversion polymorphism (Fig. 3D,G). Through IGC and inversions, *B7* sequence can occupy any of the four canonical *NPIP* locations in this cluster—thus effectively “relocating” or “repositioning” as a result of IGC. Additionally, 8/13 configurations also have a 355 kbp inversion with respect to T2T-CHM13, and only the inverted orientation configurations carry the *B6* or *B9* genes. At 99.6% sequence identity to the reference genome, the I9 inverted region is among the most divergent (top 9.5%) euchromatic regions of the human genome. We also observed 1.6 and 1.3 Mbp inversions at chromosome 16p12.3 and p12.2, respectively (Fig. 3D,E). Altogether, of the nine loci containing *NPIP* paralogs, only the locus at chromosome 16p13.3 containing *B2* is structurally invariant.

### Diversity-based tests of selection

The *NPIP* gene family members were previously shown to harbor a significant excess of amino-acid replacements, exhibiting one of the most extreme signals of positive selection in the human and African ape lineage (i.e., dN/dS>1.0) (1). To assess whether positive selection is still ongoing in the human population and narrow down signatures to individual loci, we performed complementary tests of Tajima’s D and *nS*_L_ for extended haplotype homozygosity (53, 54). Tajima’s D compares the number of segregating sites to pairwise differences to find deviations from the neutral expectation; negative values correspond to an abundance of rare alleles and are consistent with positive selection, while positive values correspond to a scarcity of rare alleles and are consistent with balancing selection. *nS*_L_ (number of segregating sites by length) is a test of extended haplotype homozygosity designed to detect recent hard and soft selective sweeps (54) and is more robust than Tajima’s D to artifactual signals arising from bottlenecks and population growth. Unlike other haplotype-based tests of selective sweeps like iHS, it is robust to phasing errors and does not rely on detailed recombination maps, as such maps are either nonexistent or unreliable in SD regions (Methods) (55).

Previous attempts have been confounded by the inability to align short reads to these duplicated regions, but our contiguous haplotype-resolved assemblies now allow us to investigate whether there is evidence of selective sweeps across these regions in the human population. We calculated *nS*_L_ and Tajima’s D with HiFi assemblies, restricting to individuals of African ancestry where we have the most samples of any individual superpopulation and to avoid bias due to the out-of-Africa bottleneck and subsequent expansions (56). To evaluate consistency of Tajima’s D within unique sequence flanking SDs, we compared Tajima’s D using short reads from the 1KG (Gambian individuals). Signals are comparable in unique regions (Fig. 4) but dropout over SDs for short-read sequence data. Using Tajima’s D, we find positive selection signatures in the 1% most extreme chromosome-wide for *NPIPB9*, *B12*, and *B15*, while *A1*, *A2*, *A5*, *B3*, *B4*, *B11*, *B13*, and *B14* are within the 5^th^ percentile. *B7* and *A7* are, however, within the 5% most extreme windows for balancing selection (Fig. 4A). Similarly, with *nS*_L_ we find signatures of selective sweeps for *B7*, *B9*, and *B15* within the 1^st^ percentile of most extreme values, and for *A8* within the top 5^th^ percentile (Fig. 4C,D). The only two regions of consecutive *nS*_L_ values in the first percentile correspond to *B7/9* and *B15 NPIP* loci. However, *B3*, *B4*, *B5*, *B7*, and *B9* are located within or near the boundaries of inversions, raising the possibility that suppressed recombination may be contributing to this signal (Fig. 3D,E).

**Figure 4.**
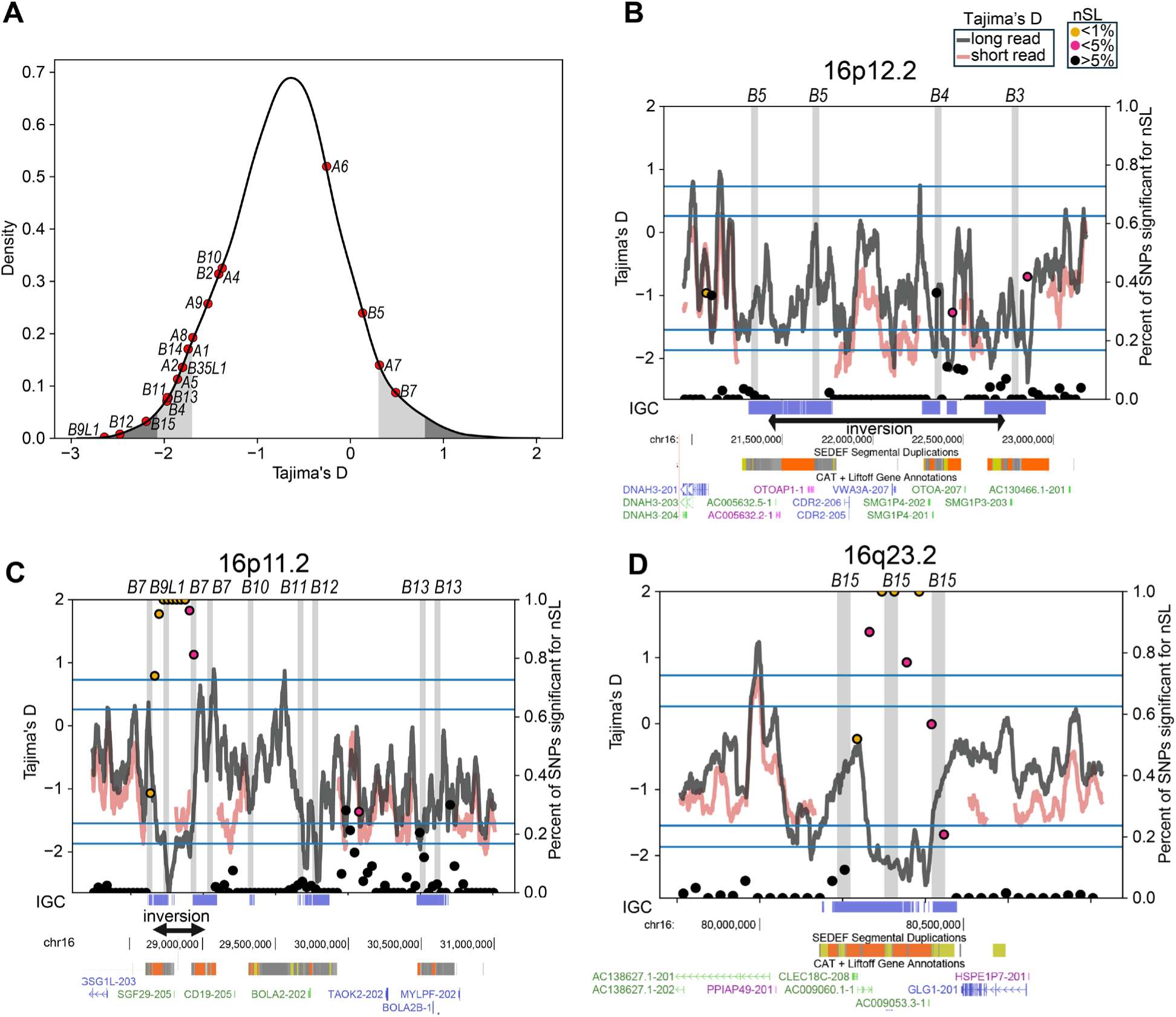
Selection signatures at *NPIP* loci in the human population. **A**) Tajima’s D distribution on chromosome 16, based on alignment of long-read sequence and assembled human haplotypes. The most extreme 1% and 5%, both positive (balancing selection) and negative (positive selection) are colored in gray and dark gray across the chromosome 16 distribution with the values for specific *NPIP* paralogs windows highlighted (red dots). **B-D**) Results of Tajima’s D and *nS*_L_ selection tests for three loci showing signatures of positive selection. Short-read (red) and long-read Tajima’s D results (gray lines) are shown. *nS*_L_ values are plotted as filled circles with color indicating significance. Known inversion polymorphisms are indicated (black arrows below) along with SDs and T2T gene annotations. Horizontal lines indicate 1% and 5% thresholds for Tajima’s D, both positive and negative. Vertical gray highlights indicate locations of *NPIP* paralogs, with gene names and SDs (SEDEF) shown below. Regions with detected IGC events are shown below in blue.

As Tajima’s D has not previously been assayable within SDs, which exhibit an increased rate of IGC and mutations in general when compared to the unique regions of the genome (46), we examined the empirical effect of IGC on Tajima’s D values. We find that the Tajima’s D distribution has an extended left tail when IGC-overlapping windows are included (Fig. S6A), with the first percentile at -2.07 as compared to -1.87 with IGC-overlapping windows excluded. High-identity *NPIP* duplications experience high rates of IGC; indeed, the nearest IGC-free windows to each *NPIP* paralog (up to 289 kbp distant) have less extreme Tajima’s D values (Fig. S6B). However, of the 11 *NPIP* paralogs within the 5% most extreme windows for positive selection, eight remain within the top 5% when IGC regions are excluded from this analysis: *A2*, *B9L1*, *B4*, *A7*, *A6*, *B14*, *B15*, and *B5*.

### NPIP gene models and differential tissue expression

Previous research demonstrated ubiquitous expression of *NPIP* paralogs in apes, as compared to the largely testis-specific expression in Old and New World monkeys, along with slightly different gene models for human *NPIPA* and *NPIPB* subfamilies (3, 57). With our more complete catalog of human *NPIP* paralogs, we sought to determine whether we could identify additional paralog-specific gene models, and if there is evidence of tissue-specific expression when considering particular *NPIP* paralogs instead of the family as a whole. Short-read RNA-seq does not align uniquely to *NPIP* paralogs due to their high sequence identity, preventing the construction of complete gene models and paralog-specific expression estimates. Instead, we used

PacBio HiFi sequencing of full-length cDNA (Iso-Seq), facilitating the unambiguous assignment of the majority of Iso-Seq reads to specific *NPIP* paralogs. To this end, we assembled a database of full-length non-chimeric (FLNC) cDNA generated from 1.4 billion Iso-Seq reads from 384 libraries, representing 101 human tissue and cell types (Supplementary Table S1). To complement this effort, we also performed hybridization capture experiments against select tissues using *NPIP-*targeting capture probes in order to enrich in *NPIP* FLNC molecules (Methods) (58). We extracted Iso-Seq reads aligning to any *NPIP* paralog, totaling 1.07 million reads with an average length of 1960 nt. To create paralog-specific gene models, we considered ORFs seen in at least five Iso-Seq molecules as valid and only display the most abundant and longest isoforms for each paralog (Fig. 5A).

**Figure 5.**
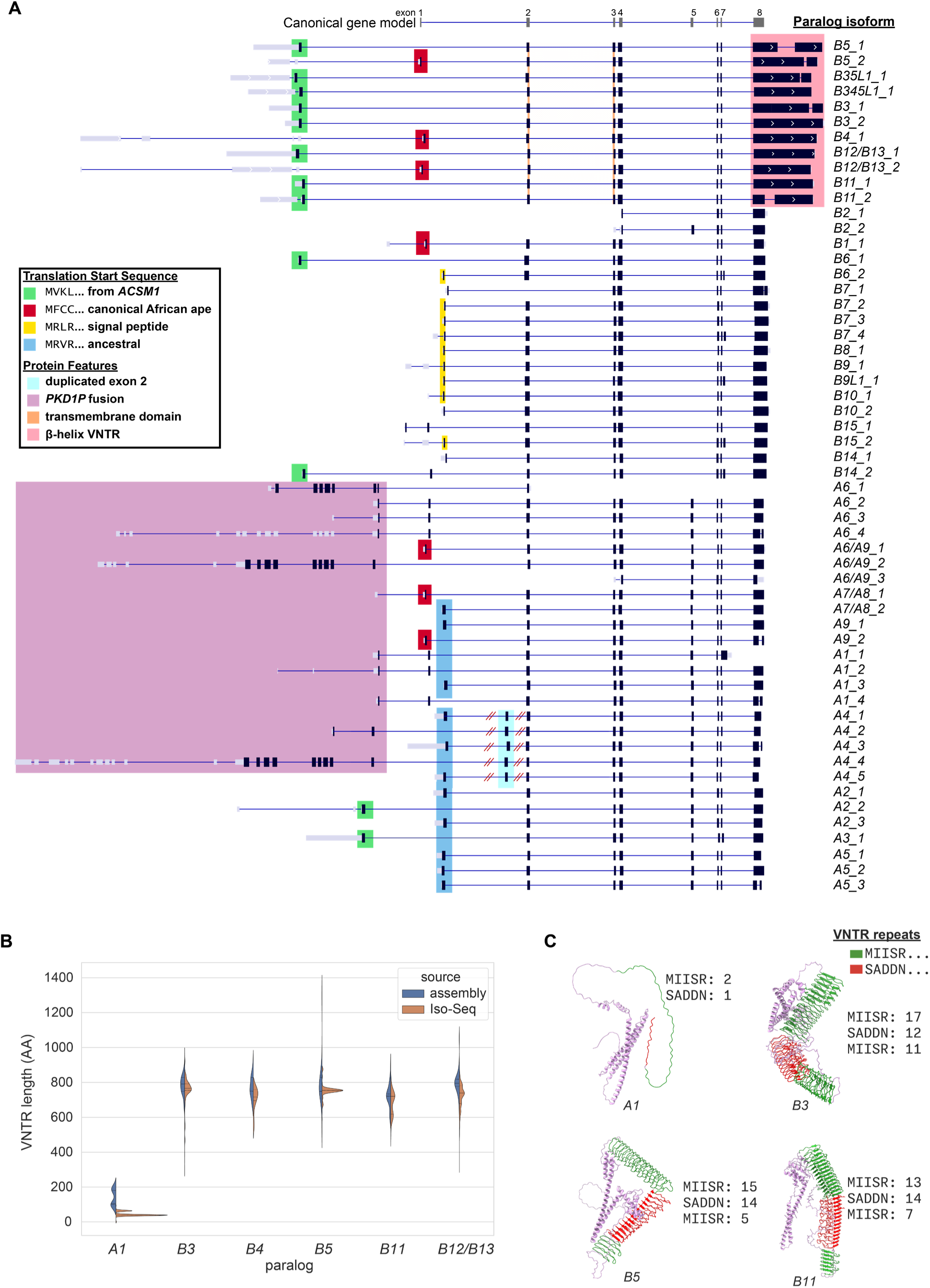
Paralog-specific gene models. **A)** Most common isoforms for each *NPIP* paralog based on full-length cDNA Iso-Seq mapping. The _x suffix indicates relative abundance (i.e., B5_1 is the most abundant B5 gene model). Predicted protein-coding regions (black), untranslated regions (gray) with different protein start sequences, and structural features are highlighted (color) over the gene models, including transcripts with the expanded protein-encoding beta-helix (pink) and the signal peptide (yellow). The canonical *NPIP* gene model is depicted (top). **B)** Comparison of VNTR length encoding the beta-helix of exon 8 in the genome assemblies versus Iso-Seq data. **C)** Predicted protein structures for four paralogs with exon 8 VNTR sizes. The copy number of repeating amino acid motifs by type are indicated and projected onto Chai-1 structure predictions. (MIISR… repeat protein domain shown in green, while frameshifted VNTR SADDN… repeat protein domain in red.)

We observe Iso-Seq molecules encoding full-length ORFs for most *NPIP* paralogs (Fig. 5). This includes four paralogs that had previously been annotated as noncoding pseudogenes, *NPIPB1P*, *NPIPP1*, *NPIPB10P*, and *NPIPB14P*, which we refer to as *B1*, *A4*, *B10*, and *B14*, respectively. The African-ape– specific *B1* paralog, the only human paralog on chromosome 18 and therefore not predisposed to the same level of structural variation, was previously reported to neither be transcribed nor maintain an ORF (3). By contrast, we find that it maintains an ORF and is expressed, albeit at low levels, in testis as well as brain organoids.

Closer inspection of these gene models reveals a considerable amount of variation in predicted amino acid composition across *NPIP* paralogs and their isoforms due to alternative promoters, differences in translation initiation, and expansion of protein-encoding variable number tandem repeats (VNTRs) at the carboxy terminus. Consequently, ORFs range in length by eightfold (155 to 1,217 amino acids). Of the 55 most common *NPIP* isoforms, only seven begin with the canonical first coding exon “MFCC…,” which is shared with African apes (3), and only 24 of these 55 were represented in RefSeq, allowing for amino acid substitutions and VNTR variation. Eleven begin with an alternate translation initiation “MVKL” sequence previously identified as the start sequence for the *NPIPB* subfamily (57). In addition to *NPIPB* paralogs, we also observe this start sequence for *NPIPA2* and *A3* and determine that this 40-amino-acid exon arose from an independent duplication of the twelfth exon of *ACSM1*, an acyl-CoA synthetase gene (Fig. S3), including half of its AMP-binding enzyme C-terminal domain (InterPro domain IPR025110). Cantsilieris et al. reported an “MRVR” start sequence in non-African ape primates, perhaps the ancestral sequence; we observe this start site used in 11 human *NPIPA* isoforms. A subset of *NPIPB* members (*B6*, *B7*, *B8*, *B9*, *B10*, *B14*, *B15*) use a previously undocumented “MRLR” start site, encoding a 19-26 amino acid signal peptide, as predicted by SignalP-6.0 (59). Though the sequence that encodes the signal peptide is present in all human *NPIP* paralogs and shared with nonhuman primates, we estimate the clade that uses this sequence as its transcription start site to be human specific, arising ∼2.6 mya during the evolution of our lineage.

Finally, for the six *NPIP* paralogs adjacent to *PKD1* pseudogenes (*A1*, *A4*, *A6*, *A7*, *A8*, *A9*), we observe 10 distinct *PKD1-NPIP* fusions, four of which are multi-exonic linking up to nine *PKD1* exons (530 aa) with eight *NPIP* exons (343 aa). Remarkably, these fusions maintain long *PKD1-NIP* ORFs up to 843 aa in length. *PKD1* variants are implicated in polycystic kidney disease, as well as estimated glomerular filtration rate (60). Though the *NPIP*-adjacent *PKD1* copies have been considered pseudogenes because the *PKD1* duplications are truncated and do not encode full-length genes (61), there is precedent for truncated genes to be functional. Several partial gene duplications like *SRGAP2C*, *NOTCH2NL*, and *ARHGAP11B* have been shown to be functional through dominant negative interaction (62–64). The role of these *PKD1-NPIP* fusion transcripts is unknown.

The final coding exon of the human-specific *NPIPB* subfamily (*B3*, *B4*, *B5*, *B11*, *B12*, *B13*) contains an expanded in-frame VNTR. Our analysis of 169 haplotypes and hundreds of cDNA libraries demonstrates that even within single paralogs, the copy number of this VNTR is variable among individuals. The VNTR encodes a repetitive amino acid motif of 19 (SADDNLKTPSERQLTPLPP) or 23 (SADDNIKTPAERLRGPLPPSAPP) residues, with the two lengths alternating. Within each paralog, the sequence frameshifts from the SADDN… form (7-15 repeats) to a MIISRHLPSVSSLPFHPQLHPQQMI form (6-14 repeats), and back to SADDN… (5-11 repeats) in the genomic annotations, resulting in a repeat domain ranging in size from 297 to 1,298 amino acids (Fig. 5B). Analyzing Iso-Seq cDNA directly, we observe 8,986 molecules sharing this VNTR switching pattern with up to 25, 20, and 17 repeat units. Computational protein structure prediction suggests that both frames of the VNTR may form a left-handed beta helix, with each VNTR unit corresponding to an additional turn of the helix, kinked as the frame shifts between these two amino acid motifs (Fig. 5C). The gene models for this subfamily also encode a transmembrane domain as predicted by DeepTMHMM (65). We estimate this specific gene and protein structure of *NPIP* is, once again, human-specific arising ∼3.1 mya (Fig. 2A, Fig. S1).

We attempted to assess paralog-specific expression levels of *NPIP* paralogs leveraging both Iso-Seq reads and short-read RNA-seq using a unique k-mer approach to specifically tag the short-read data. To determine paralog identity, the 1.07 million *NPIP* Iso-Seq reads were aligned to each of the 169 assembled haplotypes, recording the location of best mapping and requiring a difference of at least one additional mismatch to the next-best mapping paralog to consider a read uniquely identified. The approach allowed us to assign ∼55.6% of Iso-Seq reads to create paralog-specific gene models. Grouping highly similar paralogs (*A2/3*, *A6-9*, *B3-5*, and *B12/13*) allowed us to assign ∼93.2% of Iso-Seq reads for expression analysis. While these estimates are not quantitative due to errors and biases inherent in library preparation and sequencing, we observe relative and reproducible differences in paralog expression across tissue types. Comparing expression between tissue types, specific clusters of paralogs have increased relative expression in distinct tissues. In particular, *NPIPA1*, *A5*, *A6-9*, *B3-5*, and *B12/13* show increased expression in fetal or adult brain relative to other tissues, while *A2-3*, *A4*, *B1*, *B2*, *B6-9*, *B10*, *B14*, and *B15* retain the presumed ancestral testis-enriched expression pattern (Figs. 6A, S4) (57). Immune function-related tissues like tonsil, B cells, granuloma, and blood also significantly overexpress paralogs seen in brain or testis.

**Figure 6.**
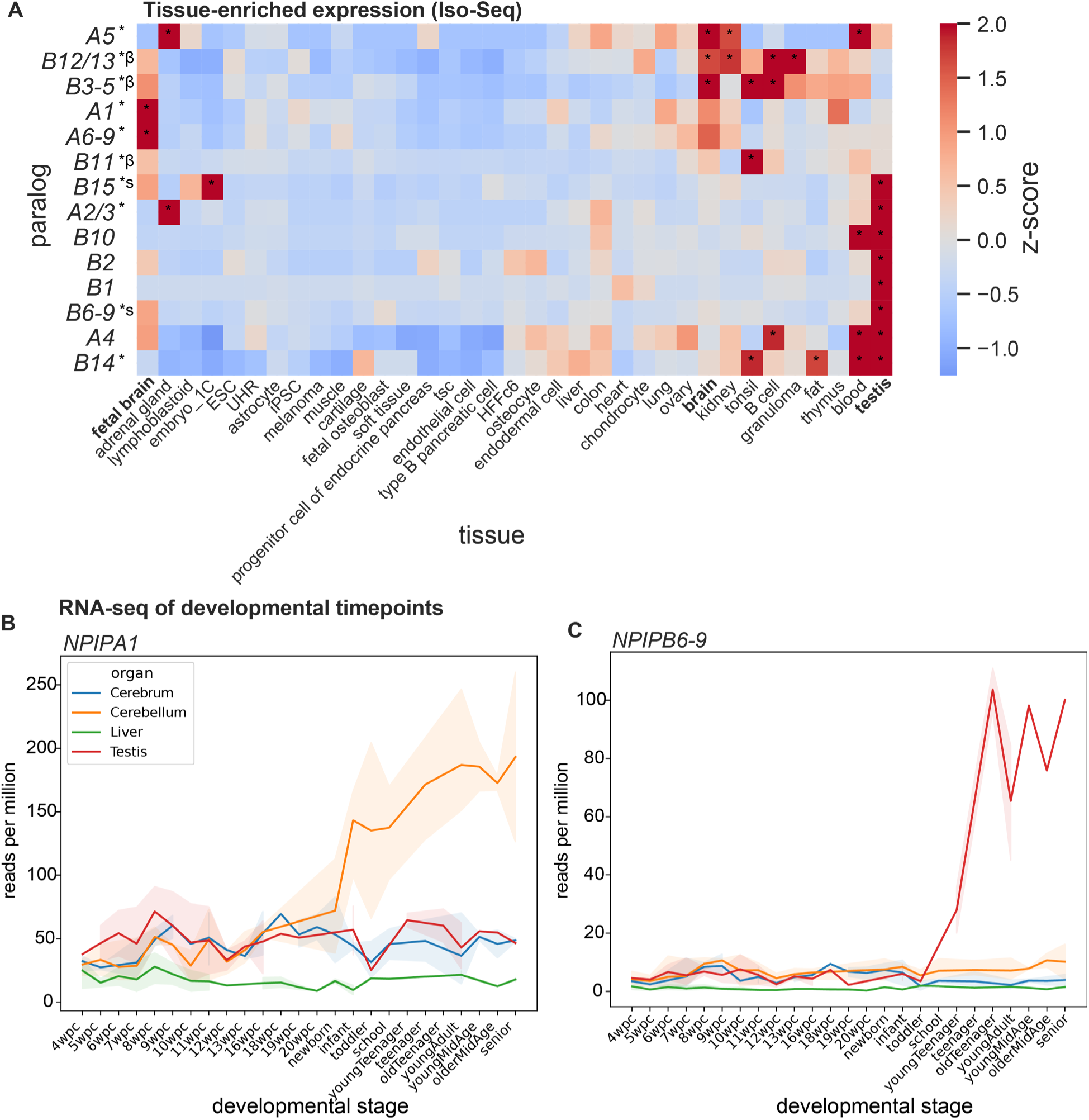
Variable expression of *NPIP* paralogs across tissues, cell types, and developmental timepoints. **A)** Relative enrichment of Iso-Seq expression estimates for 35 tissues, clustered with unweighted pair group method with arithmetic mean (UPGMA). Significantly positive z-scores are indicated with * (p<0.05). Paralogs with selection signatures are indicated with * at left; beta helix: β; signal peptide: s. **B-C)** Short-read RNA-seq expression estimates for human developmental timepoints in four tissues for *NPIPA1* and *NPIPB6-9* paralogs (aggregate), using unique k-mers for paralog identity. Transparent error bands represent 95% confidence interval of replicates.

Finally, we also attempted to define developmental time-point specificity by classifying short-read RNA-seq reads from an atlas of organ development (66) based on the presence of uniquely identifying k-mers from the 169 haplotypes. Paralogs that contained few uniquely identifying k-mers were combined into larger paralog groups for this analysis (Methods). Altogether, 25.2% of reads containing any *NPIP* k-mer (n=1.06 million) were uniquely assigned to a paralog or paralog group. *NPIPA1*, *A4*, and *B3-5* tend to increase in expression in the cerebellum after birth (Fig. 6B). In contrast, *B1*, *B2*, *B6-9*, *B10*, and *B15* expression is almost entirely testis-specific with levels increasing after puberty (Figs. 6C, S5).

## DISCUSSION

The expansion of *NPIP* and its associated SDs across the short arm of chromosome 16 predispose humans to frequent recurrent pathogenic duplications and deletions associated with autism, developmental delay, epilepsy, and obesity (6–10, 67–82). Despite this negative effect on fitness, the duplications not only persist but have expanded among the African great apes, albeit most often at non-orthologous locations (2, 3). Moreover, these same sites have homogenized via IGC, ensuring high sequence identity is maintained and driving high rates of non-allelic homologous recombination associated with disease. In light of the strong signals of positive selection for this hominid gene family (1, 3, 38), we hypothesized that an evolutionary tradeoff exists between disease susceptibility and, as of yet, unknown adaptive function. In this work, we catalog using a pangenomic approach normal human variation at each *NPIP* locus and identify paralog-specific features potentially relevant to understanding the function of this enigmatic and dynamic gene family.

Because functional human-specific duplicate genes have been shown to be frequently invariant (62, 63, 83), we systematically assessed copy number for each paralog. Based on our 28 distinct human phylogenetic groups, we find only three loci that are copy number invariant (*NPIPB2*, *B11*, and *B14*). We also distinguish loci that always have at least one copy in humans although often more (*A2*, *A4*, *B12/B13*, and *B15*). It should be noted that several of these copies are associated with larger scale structural changes, such as inversion polymorphism or IGC events. Based on the human genomes we surveyed here, only the *NPIPB2* locus is invariant by all analyses. While copy number polymorphic, the ancestral locus, *NPIPA1* (1–3), shows a striking asymmetry for IGC, serving only as a donor and never an acceptor of a gene conversion event. Such asymmetry may reflect either mutational bias or functional constraint, pinpointing copies most likely to confer function.

We previously identified extreme evolutionary signatures of positive selection for *NPIP* between African ape species, in particularly the *NPIPB* subfamily, based solely on tests for an excess of amino-acid replacement among paralogs (1, 3). With highly contiguous haplotype-resolved assemblies, we were able to apply population-level selection tests for the first time. Using Tajima’s D and *nS*_L_, we find that *NPIPB9* and *B15* are within the top percentile of most extreme values for both tests on chromosome 16, while nine additional paralogs occur within the top 5^th^ percentile for at least one test. We also find some evidence of balancing selection for a few loci (*A7* and *B7*). While these findings strongly suggest ongoing positive selection in humans, caution must be exercised given the large-scale structural changes and IGC associated with these regions. While specific copies dropout if we exclude regions of IGC, we continue to observe signals of positive selection (e.g., *NPIPB15*) in contemporary human populations though we no longer find evidence of balancing selection (Fig. S6). Additionally, *NPIPB3-B9* are located within or near the boundaries of inversions, raising the possibility that suppressed recombination may be contributing to this signal, including extended haplotypes (Fig. 3D-E). These signals may not, however, be mutually exclusive with inversions enriched for adaptively evolving genes (84, 85). Such is the case of the 17q21.31 inversion polymorphism—a locus associated within increased fecundity (49, 86), positive selection in humans (87, 88), and the dynamic evolution of newly minted gene family *LRRC37A1/2* (89–91) expressed highly in human astrocytes (92).

With a database of 1.4 billion FLNC reads (Iso-Seq), we were able to comprehensively construct paralog-specific gene models, of which 56% (31/55 most abundant isoforms) had not been previously described in RefSeq. All but one paralog maintains a full-length ORF, while *B2*, the most copy number invariant, is predicted to encode a truncated protein. The full-length gene models reveal new features—such as *NPIP* subfamilies gaining a start sequence co-opted from *ACSM1*, a novel signal peptide, transmembrane domain, or a variably sized coding VNTR that is predicted to form a beta helix and, therefore, alter the protein structure. We estimate that the signal peptide and beta helix evolved independently 2.6-3.1 mya and are innovations specific to the human lineage of evolution. The specificity afforded by long-read sequencing or paralog-specific k-mer analysis also reveals tissue-specific differences. For example, the subfamily encoding the novel signal peptide includes the two paralogs with the strongest signal of positive selection (*B6-9* and *B15*). This set shows testis-enriched expression, and analysis of a short-read development dataset additionally indicates that these paralogs increase in abundance at the onset of puberty. The paralogs with the novel beta helix and transmembrane domain, by contrast, tend to be enriched in brain samples (*B3-5*, *B12/B13*), with *B12* also among the strongest signals of positive selection.

In summary, the dynamic changes in copy number, gene model, and expression specificity across *NPIP* paralogs, along with strong signals of positive selection, suggest neofunctionalization of specific copies during human evolution. Both the paralogs that have maintained the presumed ancestral testis expression pattern (*B15*) and those that have gained enriched brain expression (*B12/B13*) exhibit clear signatures of positive selection. Concurrently, these two subfamilies evolved radically distinct gene models and associated protein structural changes in the human lineage. All of these changes have occurred and potentially been accelerated in a milieu of recurrent structural variation and IGC. Notwithstanding, it is noteworthy that *B15* and *B12/B13* are among just four paralogs that are sometimes duplicated but never deleted among humans, a potential indication of their intolerance to loss. Now that the organization and variation of these oft-overlooked loci has been resolved and their variation and gene structures understood as part of human pangenomic efforts, the next step will be associating this variation with human phenotypes, including disease.

## METHODS

### Short-read copy number estimation

We applied fastCN to high-coverage Illumina data for 2,609 unrelated individuals from the 1KG to estimate *NPIP* copy number (36, 93, 94). Short-read shotgun sequences from each individual are split into 36 bp segments and aligned to a reference genome (up to two single-nucleotide mismatches) allowing copy number to be estimated (94). Windows overlapping the exon 8 VNTR were excluded from copy number estimation to avoid biasing the estimate.

### *NPIP* gene identification

To identify *NPIP* gene locations within assemblies, we aligned the ancestral *NPIP* locus from GRCh38 (chr16:14,935,711-14,954,790) to each haplotype separately with wfmash (v0.7; parameters: -p 80 --num-mappings-for-segment=10000) and minimap2 (v2.22; parameters: -x map-ont -f 5000 -N 300 -p 0.5) (95, 96), restricting to aligned regions of at least 15 kbp. We also applied DupMasker (v1.11) (97) to identify the LCR16a duplicon where *NPIP* is located (SD9443). DupMasker identified additional copies of *NPIP* only in nonhuman primates but was not necessary for detecting *NPIP* copies in human haplotypes.

### Gene annotation

We annotated genes on each haplotype with Liftoff (v1.6.3; parameters: -flank 0.1 - polish -sc 0.85 -copies -mm2_options=“-a --end-bonus 5 --eqx -N 10000 -p 0.3 -f 1000”) (98), using protein-coding genes in GENCODE v44 (99) on GRCh38 as the reference annotation set.

### Phylogenetic paralog identity

We created an MSA of *NPIP* genes from each human and *Pongo pygmaeus* haplotype using MAFFT (v7.487; FFT-NS-2) (19). To create a phylogenetic tree of *NPIP* paralogs, we trimmed VNTRs, exons, and poorly aligned regions from the MSA visually. We estimated a maximum-likelihood phylogeny from this MSA using IQ-TREE (v2.2.3 COVID-edition; parameters -B 1000 -alrt 1000) with the GTR+F+R6 substitution model selected with ModelFinder (100, 101). Ultrafast bootstrap and SH-aLRT were used as measures of clade confidence (102, 103). Clades were named based on annotations of GRCh38, T2T-CHM13 v2.0, and T2T-HG002 and defined with SH-aLRT branch support values >75.

### Assembly validation

Assembled regions were validated with NucFreq, Flagger, and GAVISUNK depending on availability of orthogonal sequencing data (15, 39, 45, 104). Regions with no read support, only ONT support, or only HiFi support were removed. Flagger and NucFreq were applied to hifiasm and Verkko assemblies and excluded erroneous, falsely duplicated, collapsed, low confidence, or unreliable blocks. GAVISUNK was applied to hifiasm assemblies as described in Vollger, 2023, and supported regions were kept for downstream analyses (46). Only assemblies that were contiguous between proximal and distal non-segmentally duplicated marker genes were considered.

### Locus configuration comparisons

To compare structural configurations for each *NPIP* locus across samples, 10 loci were defined based on adjacent non-duplicated genes from Liftoff annotations. Configurations were defined based on order and orientation of *NPIP* paralogs and protein-coding genes from the Liftoff annotations relative to adjacent marker genes. Only configurations that passed assembly validation in at least one haplotype and were detected in at least two haplotypes were considered for further analysis. To calculate rearrangement distance between each configuration at each locus, the order and orientation of DupMasker annotations of at least 1 kbp and protein-coding marker genes were used as input to the capping-free double-cut-and-join indel model (52), and the matrix of pairwise rearrangement distances was transformed into a midpoint-rooted neighbor-joining tree with Bio.Phylo (105). The resulting trees, gene annotations, and DupMasker content were visualized with custom scripts and Baltic (106).

### VNTR analysis

To measure the length of *NPIP* exon 8 VNTRs, Tandem Repeats Finder (TRF v4.10; parameters 2 5 7 80 10 10 2000 -d -ngs) was applied to each *NPIP* copy from each haplotype (107). The longest contiguous region of tandem repeats with period of at least 40 bp was considered for each *NPIP* copy. Exon 8 VNTR size was also called directly from Iso-Seq predicted ORFs by counting substrings containing “SADD” and “ISR” for the two frames of the repeat.

### Gene model and ORF prediction

Iso-Seq reads were used to generate gene models on each human haplotype with PacBio Pigeon and SQANTI3 (v5.2), and ORF sequences with GeneMark (108, 109). Only uniquely-mapping Iso-Seq reads were used for gene model prediction, defined as a delta of at least one additional mismatch between the best-mapping paralog and second-best mapping. Mono-exonic reads were excluded. For comparison to gene models, ORFs were called directly from each *NPIP* Iso-Seq read with ANGEL (110), keeping the longest ORF per molecule.

### Selection analyses

PAV v2.4.0.1 was used to call variants for each assembled HGSVC3 (Freeze 4) haplotype relative to T2T-CHM13 v2.0 (14). Analysis was restricted to chromosome 16, containing all but one *NPIP* paralog, and African samples (n=20) to reduce the impact of population bottlenecks in the human demography. Variants were restricted to biallelic SNPs with BCFtools (111). Tajima’s D was estimated for sliding 30 kbp windows with VCF-kit (112). For comparison, Tajima’s D was called in the same way using high-coverage Illumina data for Gambian samples in the 1KG samples (n=119), restricting to 95% mappable regions as defined by the Genome in a Bottle Consortium (113). *nS*_L_ was called for 30 kbp windows with selscan v2.0.2 (55), for PAV (long-read) and 1KG (short-read) samples. PAV African *nS*_L_ results were jointly normalized for variant frequency with 95% mappable short-read calls for Gambian (GWD) samples (parameters --nsl --bins 100 --qbins 10 --min-snps 10 --bp-win --winsize 30000). Windows overlapping a T2T-CHM13 v2.0 *NPIP* copy by at least 5 kbp were considered valid.

### Protein structure prediction

For the long exon 8 VNTR isoforms predicted with SQANTI3, protein structures were predicted with Chai-1, using MSA-free mode as *NPIP* does not have the deep homology exploited by MSA-based methods for structure prediction (114). Protein structure predictions were visualized with ChimeraX (115).

### Short-read RNA-seq expression analysis

To quantify *NPIP* paralog-specific expression from short-read RNA-seq, reads were first aligned to T2T-CHM13 v2.0 with hisat2 (116). Jellyfish was used to find all possible 31-mers from *NPIP* gene models that were not found in the rest of the T2T-CHM13 v2.0 genome (117). The uniqueness of each k-mer was classified by the number of T2T-CHM13 *NPIP* paralogs in which it was found, and paralogs were iteratively merged to form detectable paralog groups until each group contained at least five uniquely identifying k-mer positions. A custom script was then used to count each identifying k-mer with each RNA-seq read and classify reads by paralog group.

### Visualization

Structural variant and phylogenetic visualizations were created with SVbyEye, archaeopteryx, augur, MEGA, and augur/auspice (118–121).

### Timetree analysis

A timetree was inferred by applying the LSD2 method (41) to a maximum likelihood neutral *NPIP* phylogenetic tree estimated with IQ-TREE (100), including a single human sequence for each paralog extracted from CHM13, CHM1, GRCh38, HG002, or PNG15, along with ape sequences from the primary *Pan troglodytes*, *Pan paniscus*, *Gorilla gorilla*, *Pongo pygmaeus*, *Pongo abelii*, and *Symphalangus syndactylus* haplotypes from the T2T Primate Project (v2.0) (42), and the ancestral *NPIP* gene from *Macaca fascicularis* (43) as the outgroup, aligned with MAFFT (19). The timetree was calibrated with the *Homo*-*Macaca* divergence set as 28.8 mya. The estimated log likelihood value of the tree is -91,940.323. There were 11,202 total positions in the final dataset.

### Probe design, cDNA generation, enrichment, and sequencing

Biotinylated oligonucleotide probes targeting *NPIP* (Supplementary Table S3) were designed to enrich in *NPIP* FLNC cDNA as described in Dougherty, 2018. Briefly, probes were designed to target constitutive exons for subfamilies A and B (exons 2, 3, 5, 6, and 7), avoiding repeat-masked sequence. Then, 5’ biotinylated sense strand oligonucleotides were synthesized by IDT for *NPIP* enrichment.

cDNA were generated using the Clontech SMARTer PCR cDNA Synthesis Kit for CHM1 (BioSample SAMN02205338), adult brain (Clontech catalog no. 636102), fetal brain (Clontech catalog no. 636106), heart (Takara catalog no. 636532, lot 1902102A), lung (Origene sample ID FR5B3386C1), ovary (Origene sample ID FR00027E9B), thymus (Origene sample ID FR5B338054), and testis (Takara catalog no. 636533 lot 1402004; BioSample SAMN15935045). For fetal brain and testis samples, cDNA were also generated using the TeloPrime Full-Length cDNA Amplification Kit V2 (Lexogen), which aims to avoid generating truncating cDNA by requiring the 5’ mRNA cap.

Unenriched polyA cDNA were sequenced from heart, lung, ovary, and thymus samples, which were barcoded and pooled on a PacBio Sequel II SMRT cell with 30-hour movie time and two-hour pre-extension.

Hybridization capture was performed on cDNA from the remaining tissues using the biotinylated *NPIP* probes, as described in Dougherty, 2018. A single Sequel SMRT cell was used for each of CHM1 and adult brain, with the remaining samples barcoded and pooled for sequencing.

Iso-Seq data from previous publications and public data depositions were obtained from ANVIL, ENCODE, and SRA, as referenced in Supplementary Table S1, and analyzed together with our generated FLNC data (21–35).

## ACKNOWLEDGMENTS

This work was supported, in part, by US National Institutes of Health (NIH) grants R01HG002385 to E.E.E. and 5R00HG011041 to P.H. E.E.E. is an investigator of the Howard Hughes Medical Institute.

This article is subject to HHMI’s Open Access to Publications policy. HHMI lab heads have previously granted a nonexclusive CC BY 4.0 license to the public and a sublicensable license to HHMI in their research articles. Pursuant to those licenses, the author-accepted manuscript of this article can be made freely available under a CC BY 4.0 license immediately upon publication.

## SUPPLEMENTARY FIGURES

**Figure S1.**
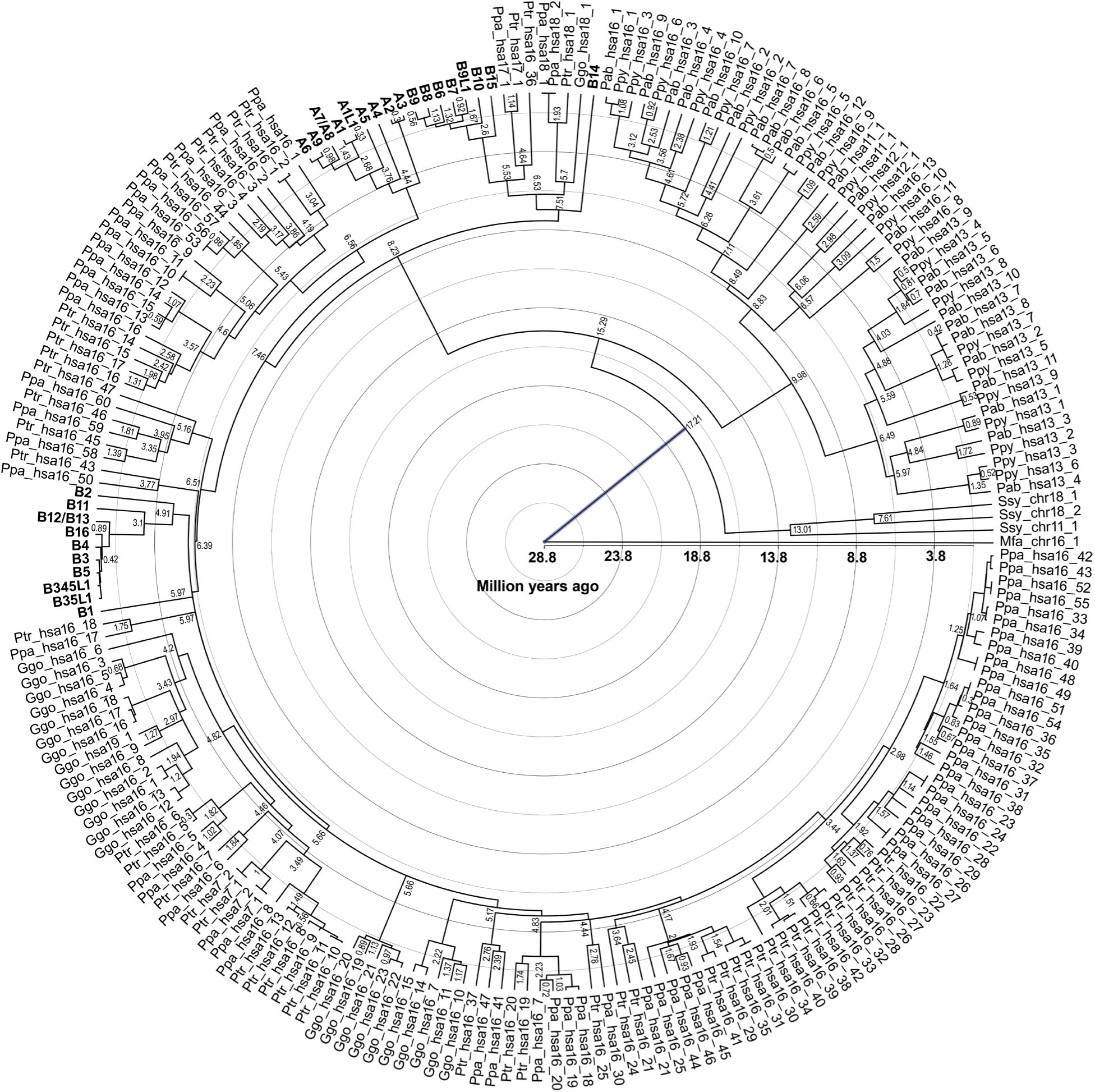
Timetree of human and ape *NPIP* duplications. The estimated age of *NPIP* paralogs for humans and ape species *Pan troglodytes* (Ptr), *Pan paniscus* (Ppa), *Gorilla gorilla* (Ggo), *Pongo pygmaeus* (Ppy), *Pongo abelii* (Pab), *Symphalangus syndactylus* (Ssy) is shown on a neutral phylogeny. The tree is rooted to the single copy ancestral *NPIP* from *Macaca fascicularis* (Mfa), with divergence time set to 28.8 mya. Human paralogs are bolded, and NHP paralogs are labeled by species abbreviation, chromosome number (hsa: human homologous chromosome, chr: species chromosome name), and position within the chromosome. Branch time estimates are indicated at the branch point.

**Figure S2.**
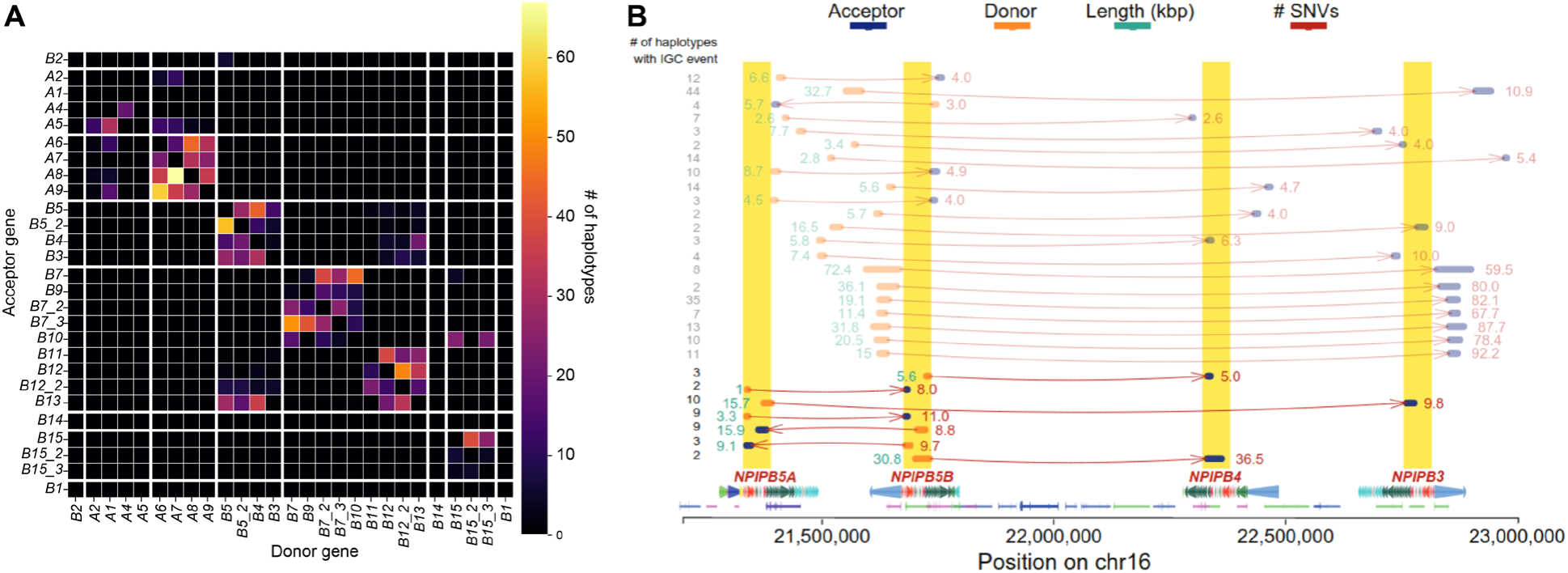
*NPIP* interlocus gene conversion (IGC) and structural changes. **A)** Counts of IGC events detected between paralogs for a subset of 94 haplotypes. X-axis labels indicate the IGC donor, and the y-axis denotes the acceptor. **B)** IGC events at the *B3-5* locus. Arrows indicate acceptors (blue) and donors (orange). The length of each event and number of SNVs are shown to the left and right, respectively.

**Figure S3.**
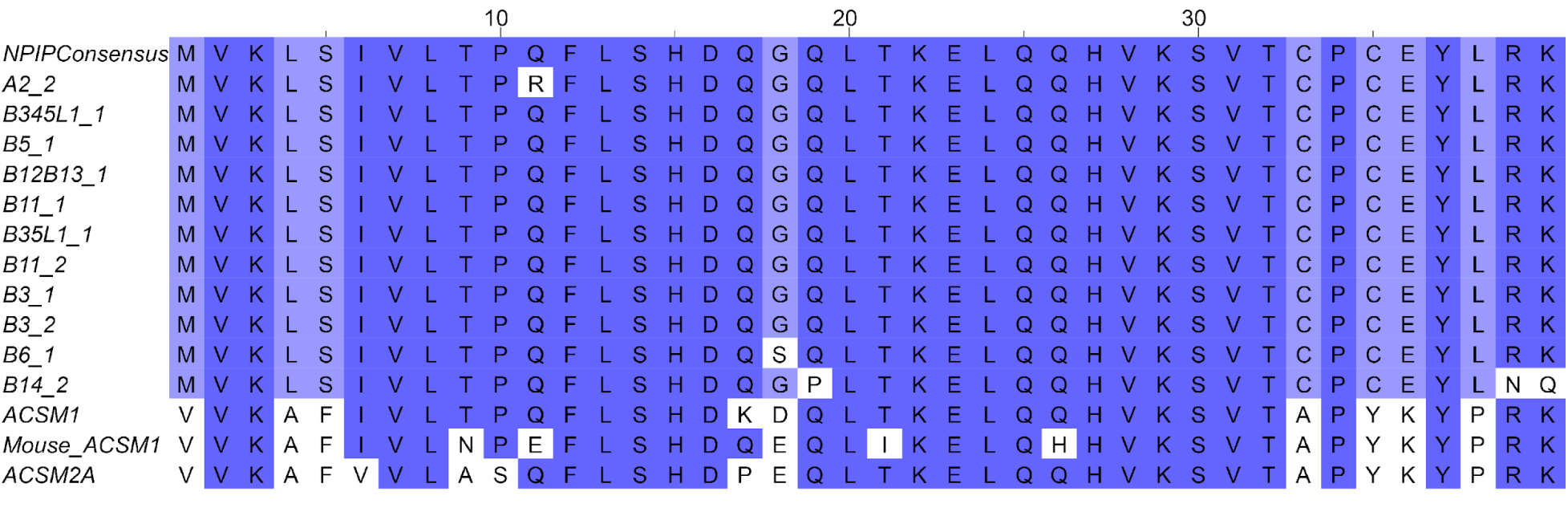
*NPIP* start sequence derived from *ACSM1*. Multiple sequence alignment of human *NPIP* paralog start sequences compared to human *ACSM1* and *ACSM2A* and mouse *ACSM1*.

**Figure S4.**
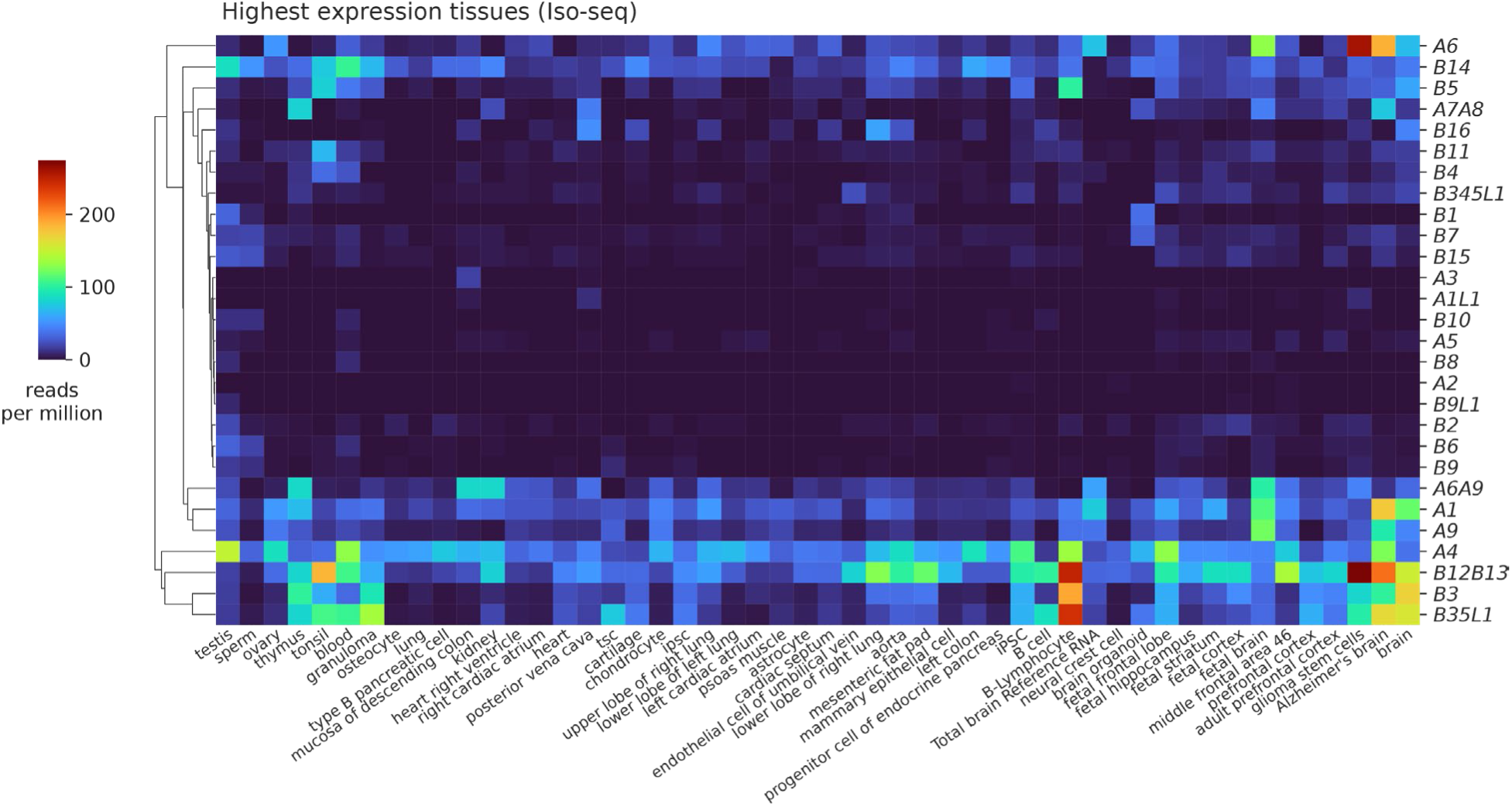
Variable expression of *NPIP* paralogs across tissues and cell types. Iso-Seq expression estimates for 50 tissues showing the highest *NPIP* expression, clustered with UPGMA.

**Figure S5.**
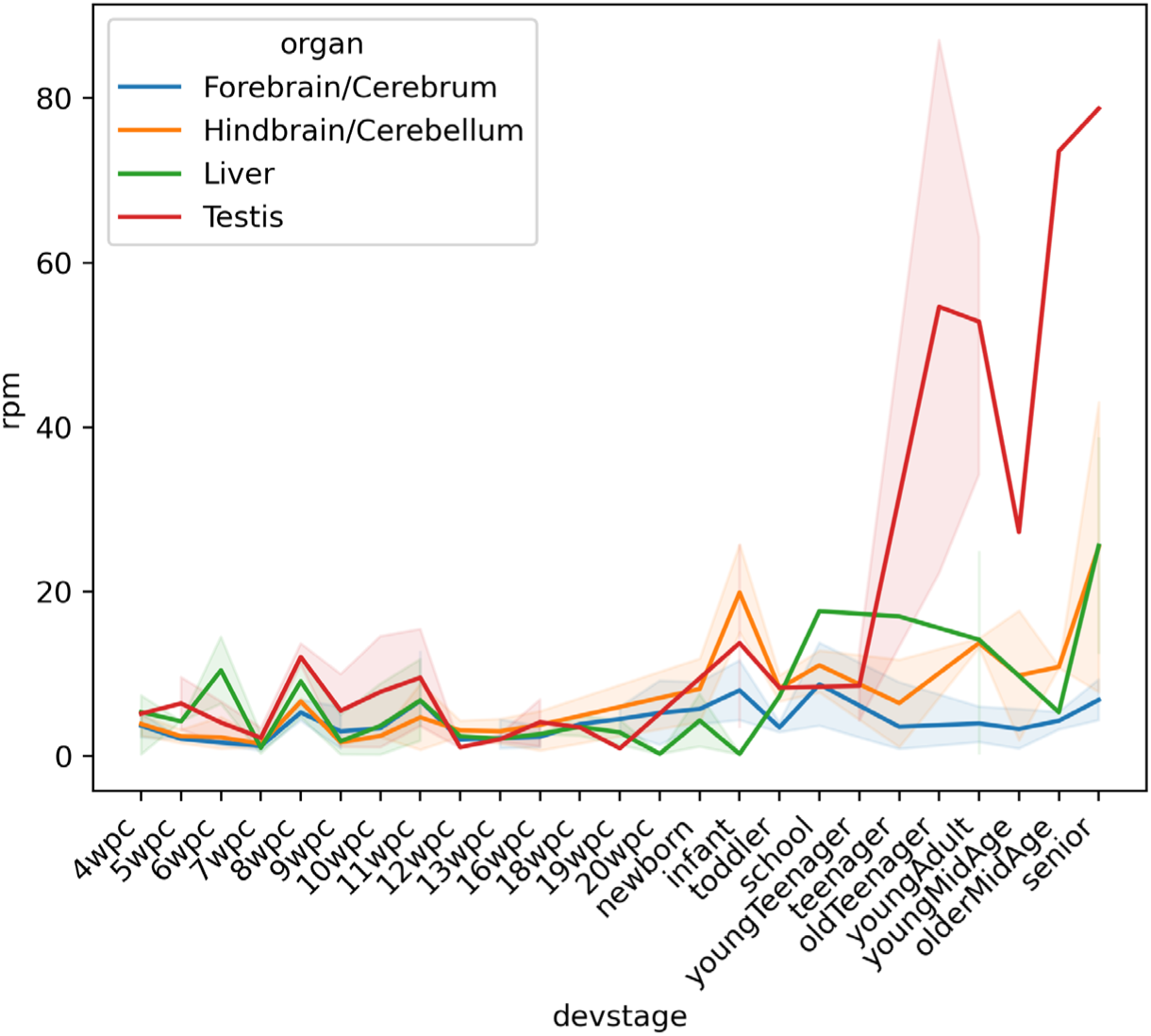
*NPIPB15* expression across development. Short-read expression estimates for human developmental timepoints in four tissues, using unique k-mers for paralog identity. Transparent error bands represent 95% confidence interval of replicates.

**Figure S6.**
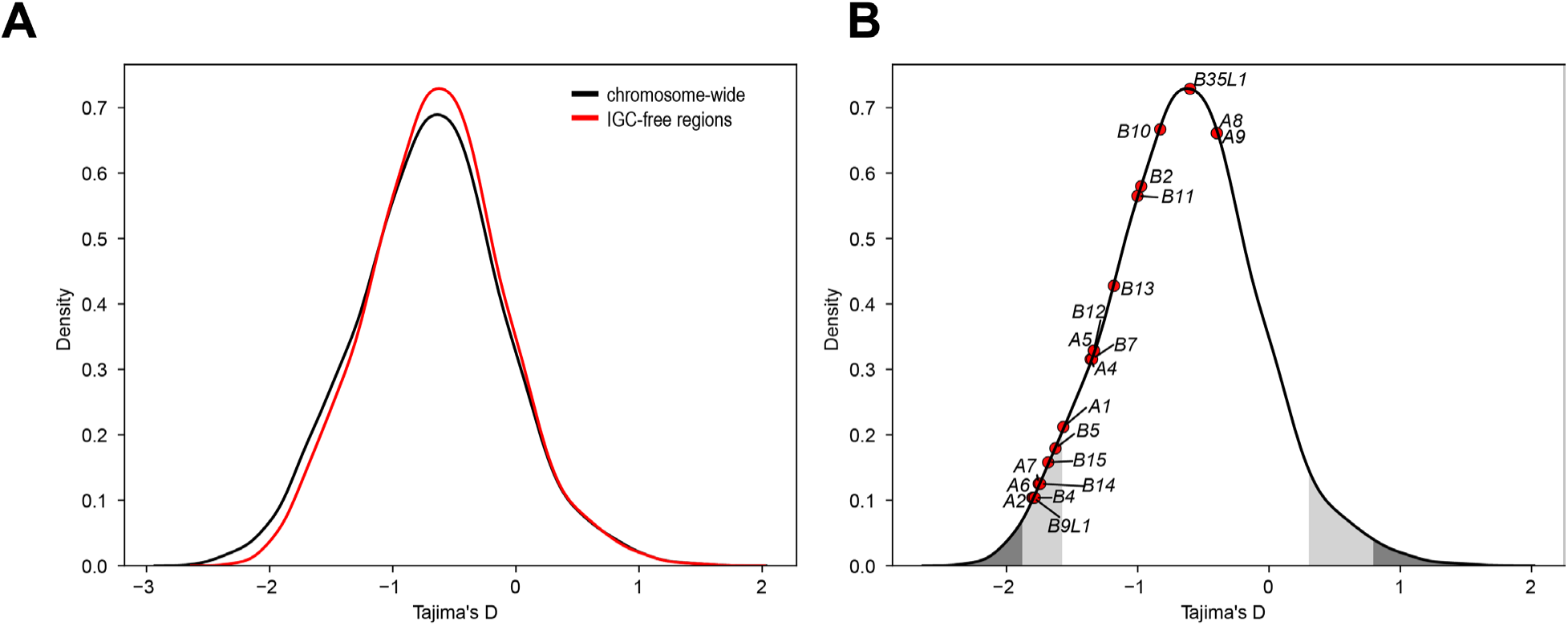
Tajima’s D distribution in IGC-free regions. **A)** Tajima’s D values for African individuals for the entirety of callable regions on chromosome 16 compared to IGC-free regions of chromosome 16. **B)** Tajima’s D values for IGC-free windows nearest to each *NPIP* paralog. The most extreme 1% and 5%, both positive (balancing selection) and negative (positive selection) are colored in gray and dark gray.

## REFERENCES

1. M. E. Johnson, et al., Positive selection of a gene family during the emergence of humans and African apes. Nature 413, 514–519 (2001).

2. M. E. Johnson, et al., Recurrent duplication-driven transposition of DNA during hominoid evolution. Proc. Natl. Acad. Sci. U.S.A. 103, 17626–17631 (2006).

3. S. Cantsilieris, et al., An evolutionary driver of interspersed segmental duplications in primates. Genome Biol 21, 202 (2020).

4. B. J. Loftus, et al., Genome Duplications and Other Features in 12 Mb of DNA Sequence from Human Chromosome 16p and 16q. Genomics 60, 295–308 (1999).

5. R. L. Stallings, S. A. Whitmore, N. A. Doggett, D. F. Callen, Refined physical mapping of chromosome 16-specific low-abundance repetitive DNA sequences. Cytogenetics and Cell Genetics 63, 97–101 (2008).

6. S. Girirajan, et al., A recurrent 16p12.1 microdeletion supports a two-hit model for severe developmental delay. Nat Genet 42, 203–209 (2010).

7. A. J. Sharp, et al., Discovery of previously unidentified genomic disorders from the duplication architecture of the human genome. Nat Genet 38, 1038–1042 (2006).

8. B. C. Ballif, et al., Discovery of a previously unrecognized microdeletion syndrome of 16p11.2– p12.2. Nat Genet 39, 1071–1073 (2007).

9. L. A. Weiss, et al., Association between Microdeletion and Microduplication at 16p11.2 and Autism. New England Journal of Medicine 358, 667–675 (2008).

10. R. A. Kumar, et al., Recurrent 16p11.2 microdeletions in autism. Human Molecular Genetics 17, 628–638 (2008).

11. Z. Jiang, et al., Ancestral reconstruction of segmental duplications reveals punctuated cores of human genome evolution. Nat Genet 39, 1361–1368 (2007).

12. P. H. Sudmant, et al., Diversity of human copy number variation and multicopy genes. Science 330, 641–646 (2010).

13. S. Nurk, et al., The complete sequence of a human genome. Science 376, 44–53 (2022).

14. P. Ebert, et al., Haplotype-resolved diverse human genomes and integrated analysis of structural variation. Science 372, eabf7117 (2021).

15. W.-W. Liao, et al., A draft human pangenome reference. Nature 617, 312–324 (2023).

16. G. A. Logsdon, et al., Complex genetic variation in nearly complete human genomes. [Preprint] (2024). Available at: https://www.biorxiv.org/content/10.1101/2024.09.24.614721v1 [Accessed 3 February 2025].

17. X. Guitart, et al., Independent expansion, selection and hypervariability of the TBC1D3 gene family in humans. Genome Res gr.279299.124 (2024). 10.1101/gr.279299.124.

18. P. Hallast, et al., Assembly of 43 human Y chromosomes reveals extensive complexity and variation. Nature 621, 355–364 (2023).

19. K. Katoh, D. M. Standley, MAFFT Multiple Sequence Alignment Software Version 7: Improvements in Performance and Usability. Mol Biol Evol 30, 772–780 (2013).

20. L.-T. Nguyen, H. A. Schmidt, A. von Haeseler, B. Q. Minh, IQ-TREE: A Fast and Effective Stochastic Algorithm for Estimating Maximum-Likelihood Phylogenies. Molecular Biology and Evolution 32, 268–274 (2015).

21. S.-J. Zhang, et al., Evolutionary interrogation of human biology in well-annotated genomic framework of rhesus macaque. Mol Biol Evol 31, 1309–1324 (2014).

22. J. M. Zook, et al., Extensive sequencing of seven human genomes to characterize benchmark reference materials. Sci Data 3, 160025 (2016).

23. Y. H. Sun, et al., Single-molecule long-read sequencing reveals a conserved intact long RNA profile in sperm. Nat Commun 12, 1361 (2021).

24. H. Kim, et al., KOREF_S1: phased, parental trio-binned Korean reference genome using long reads and Hi-C sequencing methods. GigaScience 11, giac022 (2022).

25. M. Caballero, et al., Comprehensive analysis of DNA replication timing across 184 cell lines suggests a role for MCM10 in replication timing regulation. Hum Mol Genet 31, 2899–2917 (2022).

26. A. R. Miller, et al., Pacific Biosciences Fusion and Long Isoform Pipeline for Cancer Transcriptome–Based Resolution of Isoform Complexity. The Journal of Molecular Diagnostics 24, 1292–1306 (2022).

27. F. Reese, et al., The ENCODE4 long-read RNA-seq collection reveals distinct classes of transcript structure diversity. [Preprint] (2023). Available at: https://www.biorxiv.org/content/10.1101/2023.05.15.540865v1 [Accessed 15 October 2024].

28. A. Abood, et al., Long-read proteogenomics to connect disease-associated sQTLs to the protein isoform effectors of disease. Am J Hum Genet 111, 1914–1931 (2024).

29. W. A. Cheung, et al., Direct haplotype-resolved 5-base HiFi sequencing for genome-wide profiling of hypermethylation outliers in a rare disease cohort. Nat Commun 14, 3090 (2023).

30. A. Rybak-Wolf, et al., Modelling viral encephalitis caused by herpes simplex virus 1 infection in cerebral organoids. Nat Microbiol 8, 1252–1266 (2023).

31. M. D. Schertzer, et al., Cas13d-mediated isoform-specific RNA knockdown with a unified computational and experimental toolbox. [Preprint] (2023). Available at: https://www.biorxiv.org/content/10.1101/2023.09.12.557474v1 [Accessed 15 October 2024].

32. T. D, et al., Isoform-resolved transcriptome of the human preimplantation embryo. Nature communications 14 (2023).

33. R. Garza, et al., LINE-1 retrotransposons drive human neuronal transcriptome complexity and functional diversification. Sci Adv 9, eadh9543 (2023).

34. J. H. Maeng, H. J. Jang, A. Y. Du, S.-C. Tzeng, T. Wang, Using long-read CAGE sequencing to profile cryptic-promoter-derived transcripts and their contribution to the immunopeptidome. Genome Res 33, 2143–2155 (2023).

35. M. Shimada, et al., Identification of region-specific gene isoforms in the human brain using long-read transcriptome sequencing. Science Advances 10, eadj5279 (2024).

36. M. Byrska-Bishop, et al., High-coverage whole-genome sequencing of the expanded 1000 Genomes Project cohort including 602 trios. Cell 185, 3426–3440.e19 (2022).

37. M. Rautiainen, et al., Telomere-to-telomere assembly of diploid chromosomes with Verkko. Nat Biotechnol 41, 1474–1482 (2023).

38. P. Hsieh, et al., Adaptive archaic introgression of copy number variants and the discovery of previously unknown human genes. Science 366, eaax2083 (2019).

39. P. C. Dishuck, A. N. Rozanski, G. A. Logsdon, D. Porubsky, E. E. Eichler, GAVISUNK: genome assembly validation via inter-SUNK distances in Oxford Nanopore reads. Bioinformatics 39, btac714 (2022).

40. M. R. Vollger, et al., Segmental duplications and their variation in a complete human genome. Science 376, eabj6965 (2022).

41. T.-H. To, M. Jung, S. Lycett, O. Gascuel, Fast Dating Using Least-Squares Criteria and Algorithms. Systematic Biology 65, 82–97 (2016).

42. D. Yoo, et al., Complete sequencing of ape genomes. [Preprint] (2024). Available at: https://www.biorxiv.org/content/10.1101/2024.07.31.605654v2 [Accessed 1 November 2024].

43. S. Zhang, et al., Comparative genomics of macaques and integrated insights into genetic variation and population history. bioRxiv 2024.04.07.588379 (2024). 10.1101/2024.04.07.588379.

44. D. Porubsky, et al., Gaps and complex structurally variant loci in phased genome assemblies. Genome Res. 33, 496–510 (2023).

45. M. R. Vollger, et al., Long-read sequence and assembly of segmental duplications. Nat Methods 16, 88–94 (2019).

46. M. R. Vollger, et al., Increased mutation and gene conversion within human segmental duplications. Nature 617, 325–334 (2023).

47. D. Porubsky, et al., Recurrent inversion polymorphisms in humans associate with genetic instability and genomic disorders. Cell 185, 1986–2005.e26 (2022).

48. D. Porubsky, et al., Inversion polymorphism in a complete human genome assembly. Genome Biology 24, 100 (2023).

49. M. C. Zody, et al., Evolutionary toggling of the MAPT 17q21.31 inversion region. Nat Genet 40, 1076–1083 (2008).

50. J. R. González, et al., A Common 16p11.2 Inversion Underlies the Joint Susceptibility to Asthma and Obesity. American Journal of Human Genetics 94, 361 (2014).

51. J. R. González, et al., Polymorphic Inversions Underlie the Shared Genetic Susceptibility of Obesity-Related Diseases. The American Journal of Human Genetics 106, 846–858 (2020).

52. L. Bohnenkämper, Recombinations, chains and caps: resolving problems with the DCJ-indel model. Algorithms for Molecular Biology 19, 8 (2024).

53. F. Tajima, Statistical method for testing the neutral mutation hypothesis by DNA polymorphism. Genetics 123, 585–595 (1989).

54. A. Ferrer-Admetlla, M. Liang, T. Korneliussen, R. Nielsen, On Detecting Incomplete Soft or Hard Selective Sweeps Using Haplotype Structure. Molecular Biology and Evolution 31, 1275–1291 (2014).

55. Z. A. Szpiech, selscan 2.0: scanning for sweeps in unphased data. Bioinformatics 40, btae006 (2024).

56. J. C. Fay, C. I. Wu, A human population bottleneck can account for the discordance between patterns of mitochondrial versus nuclear DNA variation. Mol Biol Evol 16, 1003–1005 (1999).

57. C. Bekpen, et al., Functional Characterization of the Morpheus Gene Family. (2017). 10.1101/116087.

58. M. L. Dougherty, et al., Transcriptional fates of human-specific segmental duplications in brain. Genome Res. 28, 1566–1576 (2018).

59. F. Teufel, et al., SignalP 6.0 predicts all five types of signal peptides using protein language models. Nat Biotechnol 40, 1023–1025 (2022).

60. J. N. Hellwege, et al., Mapping eGFR loci to the renal transcriptome and phenome in the VA Million Veteran Program. Nat Commun 10, 3842 (2019).

61. N. Bogdanova, et al., Homologues to the first gene for autosomal dominant polycystic kidney disease are pseudogenes. Genomics 74, 333–341 (2001).

62. M. Y. Dennis, et al., Evolution of Human-Specific Neural *SRGAP2* Genes by Incomplete Segmental Duplication. Cell 149, 912–922 (2012).

63. M. Florio, et al., Human-specific gene ARHGAP11B promotes basal progenitor amplification and neocortex expansion. Science 347, 1465–1470 (2015).

64. I. T. Fiddes, et al., Human-Specific NOTCH2NL Genes Affect Notch Signaling and Cortical Neurogenesis. Cell 173, 1356–1369.e22 (2018).

65. J. Hallgren, et al., DeepTMHMM predicts alpha and beta transmembrane proteins using deep neural networks. [Preprint] (2022). Available at: https://www.biorxiv.org/content/10.1101/2022.04.08.487609v1 [Accessed 1 November 2024].

66. M. Cardoso-Moreira, et al., Gene expression across mammalian organ development. Nature 571, 505–509 (2019).

67. E. L. Heinzen, et al., Rare Deletions at 16p13.11 Predispose to a Diverse Spectrum of Sporadic Epilepsy Syndromes. Am J Hum Genet 86, 707–718 (2010).

68. P. N. Pop-Jordanova, T. Zorcec, E. Sukarova-Angelovska, Duplication of Chromosome 16p13.11-p12.3 with Different Expressions in the Same Family. Balkan Journal of Medical Genetics : BJMG 24, 89 (2021).

69. I. Quintela, et al., A maternally inherited 16p13.11-p12.3 duplication concomitant with a de novo SOX5 deletion in a male patient with global developmental delay, disruptive and obsessive behaviors and minor dysmorphic features. American Journal of Medical Genetics Part A 167, 1315–1322 (2015).

70. S. Loureiro, et al., Copy number variations in chromosome 16p13.11-The neurodevelopmental clinical spectrum. Current Pediatric Research (2017).

71. C. G. F. de Kovel, et al., Recurrent microdeletions at 15q11.2 and 16p13.11 predispose to idiopathic generalized epilepsies. Brain 133, 23–32 (2010).

72. S.-Q. Kuang, et al., Recurrent Chromosome 16p13.1 Duplications Are a Risk Factor for Aortic Dissections. PLoS Genetics 7, e1002118 (2011).

73. A. Ingason, et al., Copy number variations of chromosome 16p13.1 region associated with schizophrenia. Mol Psychiatry 16, 17–25 (2011).

74. A. Ramalingam, et al., 16p13.11 duplication is a risk factor for a wide spectrum of neuropsychiatric disorders. J Hum Genet 56, 541–544 (2011).

75. F. D. Hannes, et al., Recurrent reciprocal deletions and duplications of 16p13.11: the deletion is a risk factor for MR/MCA while the duplication may be a rare benign variant. J Med Genet 46, 223– 232 (2009).

76. R. Nicolle, et al., 16p13.11p11.2 triplication syndrome: a new recognizable genomic disorder characterized by optical genome mapping and whole genome sequencing. Eur J Hum Genet 30, 712–720 (2022).

77. M. N. Loviglio, et al., Chromosomal contacts connect loci associated with autism, BMI and head circumference phenotypes. Mol Psychiatry 22, 836–849 (2017).

78. J. C. K. Barber, et al., 16p11.2–p12.2 duplication syndrome; a genomic condition differentiated from euchromatic variation of 16p11.2. European Journal of Human Genetics 21, 182 (2012).

79. G. M. Cooper, et al., A copy number variation morbidity map of developmental delay. Nat Genet 43, 838–846 (2011).

80. B. P. Coe, et al., Neurodevelopmental disease genes implicated by de novo mutation and copy number variation morbidity. Nat Genet 51, 106–116 (2019).

81. F. Antonacci, et al., A large, complex structural polymorphism at 16p12.1 underlies microdeletion disease risk. Nat Genet 42, 745–750 (2010).

82. E. G. Bochukova, et al., Large, rare chromosomal deletions associated with severe early-onset obesity. Nature 463, 666–670 (2010).

83. I. T. Fiddes, A. A. Pollen, J. M. Davis, J. M. Sikela, Paired involvement of human-specific Olduvai domains and NOTCH2NL genes in human brain evolution. Human Genetics 138, 715– 721 (2019).

84. M. Kirkpatrick, N. Barton, Chromosome inversions, local adaptation and speciation. Genetics 173, 419–434 (2006).

85. B. Charlesworth, N. H. Barton, The Spread of an Inversion with Migration and Selection. Genetics 208, 377–382 (2018).

86. H. Stefansson, et al., A common inversion under selection in Europeans. Nat. Genet. 37, 129–137 (2005).

87. L. M. Boettger, R. E. Handsaker, M. C. Zody, S. A. McCarroll, Structural haplotypes and recent evolution of the human 17q21.31 region. Nat Genet 44, 881–885 (2012).

88. K. M. Steinberg, et al., Structural diversity and African origin of the 17q21.31 inversion polymorphism. Nat Genet 44, 872–880 (2012).

89. M. C. Zody, et al., DNA sequence of human chromosome 17 and analysis of rearrangement in the human lineage. Nature 440, 1045–1049 (2006).

90. C. Bekpen, I. Tastekin, P. Siswara, C. A. Akdis, E. E. Eichler, Primate segmental duplication creates novel promoters for the LRRC37 gene family within the 17q21.31 inversion polymorphism region. Genome Res 22, 1050–1058 (2012).

91. G. Giannuzzi, et al., Evolutionary dynamism of the primate LRRC37 gene family. Genome Res. 23, 46–59 (2013).

92. K. R. Bowles, et al., 17q21.31 sub-haplotypes underlying H1-associated risk for Parkinson’s disease are associated with LRRC37A/2 expression in astrocytes. Molecular Neurodegeneration 17, 48 (2022).

93. C. Alkan, et al., Personalized Copy-Number and Segmental Duplication Maps using Next-Generation Sequencing. Nat Genet 41, 1061–1067 (2009).

94. A. L. Pendleton, et al., Comparison of village dog and wolf genomes highlights the role of the neural crest in dog domestication. BMC Biol 16, 64 (2018).

95. S. Marco-Sola, et al., Optimal gap-affine alignment in O(s) space. Bioinformatics 39, btad074 (2023).

96. H. Li, Minimap2: pairwise alignment for nucleotide sequences. Bioinformatics 34, 3094–3100 (2018).

97. Z. Jiang, R. Hubley, A. Smit, E. E. Eichler, DupMasker: A tool for annotating primate segmental duplications. Genome Res. 18, 1362–1368 (2008).

98. A. Shumate, S. L. Salzberg, Liftoff: accurate mapping of gene annotations. Bioinformatics 37, 1639–1643 (2021).

99. A. Frankish, et al., GENCODE: reference annotation for the human and mouse genomes in 2023. Nucleic Acids Res 51, D942–D949 (2023).

100. B. Q. Minh, et al., IQ-TREE 2: New Models and Efficient Methods for Phylogenetic Inference in the Genomic Era. Molecular Biology and Evolution 37, 1530–1534 (2020).

101. S. Kalyaanamoorthy, B. Q. Minh, T. K. F. Wong, A. von Haeseler, L. S. Jermiin, ModelFinder: fast model selection for accurate phylogenetic estimates. Nat Methods 14, 587–589 (2017).

102. D. T. Hoang, O. Chernomor, A. von Haeseler, B. Q. Minh, L. S. Vinh, UFBoot2: Improving the Ultrafast Bootstrap Approximation. Molecular Biology and Evolution 35, 518–522 (2018).

103. M. Anisimova, M. Gil, J.-F. Dufayard, C. Dessimoz, O. Gascuel, Survey of Branch Support Methods Demonstrates Accuracy, Power, and Robustness of Fast Likelihood-based Approximation Schemes. Systematic Biology 60, 685–699 (2011).

104. A. M. M. Cartney, et al., Chasing perfection: validation and polishing strategies for telomere-to-telomere genome assemblies. Nature methods 19, 687 (2022).

105. E. Talevich, B. M. Invergo, P. J. Cock, B. A. Chapman, Bio.Phylo: A unified toolkit for processing, analyzing and visualizing phylogenetic trees in Biopython. BMC Bioinformatics 13, 209 (2012).

106. G. Dudas, evogytis/baltic. (2024). Deposited 3 October 2024.

107. G. Benson, Tandem repeats finder: a program to analyze DNA sequences. Nucleic Acids Research 27, 573–580 (1999).

108. F. J. Pardo-Palacios, et al., SQANTI3: curation of long-read transcriptomes for accurate identification of known and novel isoforms. Nat Methods 21, 793–797 (2024).

109. J. Besemer, M. Borodovsky, GeneMark: web software for gene finding in prokaryotes, eukaryotes and viruses. Nucleic Acids Res 33, W451–454 (2005).

110. PacificBiosciences/ANGEL. (2023). Deposited 28 January 2023.

111. P. Danecek, et al., Twelve years of SAMtools and BCFtools. Gigascience 10, giab008 (2021).

112. D. E. Cook, E. C. Andersen, VCF-kit: assorted utilities for the variant call format. Bioinformatics 33, 1581–1582 (2017).

113. N. Dwarshuis, et al., The GIAB genomic stratifications resource for human reference genomes. Nat Commun 15, 9029 (2024).

114. Chai Discovery, et al., Chai-1: Decoding the molecular interactions of life. [Preprint] (2024). Available at: http://biorxiv.org/lookup/doi/10.1101/2024.10.10.615955 [Accessed 30 October 2024].

115. E. F. Pettersen, et al., UCSF ChimeraX: Structure visualization for researchers, educators, and developers. Protein Sci 30, 70–82 (2021).

116. D. Kim, J. M. Paggi, C. Park, C. Bennett, S. L. Salzberg, Graph-based genome alignment and genotyping with HISAT2 and HISAT-genotype. Nat Biotechnol 37, 907–915 (2019).

117. G. Marçais, C. Kingsford, A fast, lock-free approach for efficient parallel counting of occurrences of k-mers. Bioinformatics 27, 764–770 (2011).

118. D. Porubsky, et al., SVbyEye: A visual tool to characterize structural variation among whole genome assemblies. [Preprint] (2024). Available at: https://www.biorxiv.org/content/10.1101/2024.09.11.612418v1 [Accessed 31 October 2024].

119. Archaeopteryx. Available at: http://www.phylosoft.org/archaeopteryx/ [Accessed 31 October 2024].

120. K. Tamura, G. Stecher, S. Kumar, MEGA11: Molecular Evolutionary Genetics Analysis Version 11. Molecular Biology and Evolution 38, 3022–3027 (2021).

121. J. Huddleston, et al., Augur: a bioinformatics toolkit for phylogenetic analyses of human pathogens. Journal of open source software 6, 2906 (2021).

